# IFNγ induces epigenetic programming of human T-bet^hi^ B cells and promotesTLR7/8 and IL-21 induced differentiation

**DOI:** 10.1101/557520

**Authors:** Esther Zumaquero, Sara L. Stone, Christopher D. Scharer, Scott A. Jenks, Anoma Nellore, Betty Mousseau, Antonio Rosal-Vela, Davide Botta, John E. Bradley, Wojciech Wojciechowski, Travis Ptacek, Maria I. Danila, Jeffrey C. Edberg, S. Louis Bridges, Robert P. Kimberly, W. Winn Chatham, Trenton R. Schoeb, Alexander Rosenberg, Jeremy M. Boss, Ignacio Sanz, Frances E. Lund

## Abstract

Although B cells expressing the IFNγR or the IFNγ-inducible transcription factor T-bet drive autoimmunity in Systemic Lupus Erythematosus (SLE)-prone mouse models, the role for IFNγ signaling in human antibody responses is unknown. We show that elevated levels of IFNγ in SLE patients correlate with expansion of the T-bet expressing IgD^neg^CD27^neg^CD11c^+^CXCR5^neg^ (DN2) pre-antibody secreting cell (pre-ASC) subset. We demonstrate that naïve B cells form T-bet^hi^ pre-ASCs following stimulation with either Th1 cells or with IFNγ, IL-2, anti-Ig and TLR7/8 ligand and that IL-21 dependent ASC formation is significantly enhanced by IFNγ or IFNγ-producing T cells. IFNγ promotes ASC development by synergizing with IL-2 and TLR7/8 ligands to induce genome-wide epigenetic reprogramming of B cells, which results in increased chromatin accessibility surrounding IRF4 and BLIMP1 binding motifs and epigenetic remodeling of *IL21R* and *PRDM1* loci. Finally, we show that IFNγ signals poise B cells to differentiate by increasing their responsiveness to IL-21.

## Introduction

Systemic Lupus Erythematosus (SLE) is characterized by progressive dysregulation of the innate and adaptive arms of the immune system, which ultimately leads to loss of immune tolerance in B and T lymphocytes and the production of autoantibodies (Abs) by Ab-secreting B cells (ASCs) (1). The hallmark SLE autoAbs recognize nuclear proteins and nucleic acids (2), which are also ligands for TLR7 and TLR9 that are expressed by innate immune cells and B cells (3). SLE autoAbs bound to their autoAgs form immune complexes, which are responsible for many of the clinical manifestations of SLE, particularly those associated with organ damage (2). Consistent with the important role for B cells and ASCs in SLE pathogenesis (4), the only new drug approved to treat SLE in decades, Belimumab, targets B cells.

Inflammatory cytokines and chemokines also contribute to SLE pathogenesis (5). SLE patient PBMCs often exhibit a type I interferon (IFN) transcriptional signature and systemic IFNα is elevated in many patients (6). It is less well appreciated that IFNγ is also increased in some SLE patients (7-9) and that a distinct IFNγ transcription signature can be detected in PBMCs from a portion of SLE patients (10, 11). Interestingly, elevated serum IFNγ can be observed years before IFNα or autoAbs are detected in SLE patients and much earlier than clinical disease (12, 13). Consistent with these observations, B cells from SLE patients can exhibit signs of prior IFNγ exposure. For example, CXCR3 and T-bet, two IFNγ-inducible proteins (14), are more highly expressed by circulating B cells from SLE patients compared to healthy controls (8, 15-19). Moreover, data from mouse SLE models show that clinical disease is dependent on B cell-specific expression of the IFNγR and the IFNγ-induced transcription factors STAT1 (20-22) and T-bet in some (23, 24) but not all (21, 25) models. Taken together, these data suggest that IFNγ-driven inflammation may contribute to SLE B cell-driven pathophysiology.

Two populations of circulating B cells present in SLE patients, namely the CD11c^hi^ and IgD^neg^CD27^neg^ subsets, are reported to express T-bet (18, 19). CD11c^hi^ B cells, which are called age associated B cells (ABCs) (26, 27), and the IgD^neg^CD27^neg^ double negative (B_DN_) B cells, which are often referred to as “atypical” memory B cells (28, 29), are present in low numbers in the blood or tonsils of healthy individuals (30) and are reported to be expanded in chronically infected (29, 31-33), aging (27, 34, 35) and autoimmune individuals (26, 27), including patients with SLE (28). The CD11c^hi^ population found in SLE patients is heterogeneous and contains CD11c-expressing IgD^neg^CD27^+^ switched memory (B_SW_) cells, IgD^neg^CD27^neg^ naïve (B_N_) cells and B_DN_ cells (18). The B_DN_ population is also heterogeneous and can be subdivided using CD11c and CXCR5 into DN1 (CD11c^lo^CXCR5^+^) and DN2 (CD11c^hi^CXCR5^neg^) populations (19).

Despite extensive data showing that these overlapping populations of CD11c^hi^ B cells and B_DN_ cells are expanded in a number of human diseases (36), our understanding regarding their origin and function is incomplete. Although initial studies examining B_DN_ cells from malaria or HIV-infected individuals described these B cells as anergic (31, 37-39), more recent studies reported that the CD11c-expressing IgD^neg^CD27^+^CD21^lo^ activated B_SW_ cells from influenza vaccinated humans (40) and HIV infected patients (33), as well as the CD11c^hi^ cells from SLE patients (18) and the CD11c^hi^ DN2 cells from SLE patients (19) possess phenotypic and molecular characteristics of pre-ASCs. Both the CD11c^hi^ B cells and the more narrowly defined DN2 subset from SLE patients differentiated into ASCs following stimulation (18, 19). Moreover, the T-bet^hi^ DN2 subset from SLE patients can produce autoAbs (19), suggesting that these cells can potentially contribute to disease.

Given the fact that T-bet^hi^ DN2 pre-ASCs produce autoAbs and correlate with disease severity in SLE patients (19), we set out to identify the signals that control formation of this population and their differentiation into ASCs. Consistent with the fact that SLE DN2 cells express high levels of T-bet, we show that expansion of the DN2 cells in SLE patients correlates with systemic concentrations of IFNγ and IFNγ-induced cytokines. We further demonstrate that activation of B_N_ cells with IFNγ-producing T cells or IFNγ + TLR7/8 and BCR ligands induces formation of a T-bet^hi^ pre-ASC population that is similar to the SLE T-bet^hi^ DN2 subset. Importantly, we show that IFNγ signals are not only required for formation of the pre-ASC population but also greatly augment ASC formation, at least in part by increasing IL-21R expression and responsiveness of the cells to IL-21. IFNγ appears to enhance ASC differentiation by synergizing with BCR, IL-2 and TLR7/8 signals to promote global epigenetic changes, some of which result in greatly increased chromatin accessibility surrounding binding motifs for two key ASC commitment transcription factors, BLIMP1 and IRF4. Finally, and consistent with our hypothesis that IFNγ signals poise B cells to differentiate, we identified IFNγ-dependent differentially accessible regions (DARs) within the *IL21R* and *PRDM1* (BLIMP1) loci. These DARs are also present in the SLE patient DN2 cells, suggesting that IFNγ signals might contribute to the epigenetic changes seen in the SLE T-bet^hi^ DN2 pre-ASC population and may be critical for the formation of these likely pathogenic pre-ASCs.

## RESULTS

### T-bet is highly expressed by expanded ASC precursors in SLE patients

Recent studies (18, 19) identified and characterized B cells that are expanded in a fraction of SLE patients, including an IgD^neg^CD27^neg^ B cell subset (referred to as double negative B cells or B_DN_ cells) and a CD19^hi^CD11c^hi^ subset (referred to as age associated B cells or ABCs). These two populations, which are heterogeneous and overlapping (18, 19), were reported to contain B cells expressing the IFNγ-induced transcription factor, T-bet. This was of interest to us as we (Figure 1-figure supplement 1) and others (23, 24) showed that B cell intrinsic expression of T-bet is required for the development of autoAb-mediated immunity in SLE mouse models. Consistent with this, our studies in SLE patients (19) indicated that a subpopulation of cells within the B_DN_ population, namely the CXCR5^neg^CD11c^hi^ (DN2) subset, exhibited characteristics of pre-antibody secreting cells (pre-ASCs). We further showed that expansion of this subpopulation correlated strongly with disease activity in SLE patients (19). Given these findings, we hypothesized that T-bet would be expressed by this expanded population of pre-ASCs in the SLE patients and that expansion of these T-bet expressing B cells in SLE patients would correlate with autoAb titers in these individuals. To test this hypothesis, we first measured T-bet levels in total B cells (non-ASCs), IgD^+^CD27^neg^ naïve (B_N_), IgD^neg^CD27^+^ memory (B_SW_), IgD^+^CD27^+^ unswitched memory (B_U_) and B_DN_ cell subsets isolated from peripheral blood of healthy donors (HD) and SLE patients (Fig. 1a). We observed an expansion of T-bet^hi^ B cells within the total B cell compartment as well as in all SLE B cell subsets compared to HD controls (Fig. 1b). However, T-bet^hi^ B cells were particularly prevalent within the SLE B_DN_ compartment and correlated precisely with the frequency of B_DN_ cells present in these patients (Fig. 1c). Since the B_DN_ population is heterogeneous (19) and can be subdivided into memory CXCR5^+^CD11c^lo^ B_DN_ cells (DN1 subset) and the effector CD11c^hi^CXCR5^neg^ DN2 subset (Fig. 1d), we asked whether the T-bet was specifically expressed by the DN1 or DN2 subset. We found that the T-bet^hi^ B cells were exclusively contained within the CD11c^hi^CXCR5^neg^ DN2 subpopulation (Fig. 1e). Moreover, consistent with what has been reported for the SLE DN2 cells (19), the SLE T-bet^hi^ B cells were uniformly CD19^hi^FcRL5^+^CD23^neg^ (Fig. 1f). These data therefore indicated that the T-bet^hi^ B_DN_ subset and the previously described DN2 pre-ASC subset represent equivalent populations in SLE patients. Consistent with this conclusion, two transcription factors that are required for ASC differentiation (41), Blimp1 and IRF4, are expressed at intermediate levels in T-bet^hi^ B_DN_ cells from SLE patients relative to CD27^hi^CD38^hi^ ASCs and T-bet^lo^ B cells (Fig. 1g-i). In addition, we observed a strong positive correlation between the frequency of T-bet^hi^ B_DN_ cells and anti-Smith autoAb levels in our cohort of SLE patients (Fig. 1j). Thus, the T-bet^hi^ B_DN_ subset, which we now refer to as the SLE T-bet^hi^ DN2 subset, exhibits phenotypic characteristics of pre-ASCs and are most expanded in patients with the highest autoAb titers.

**Figure 1.**
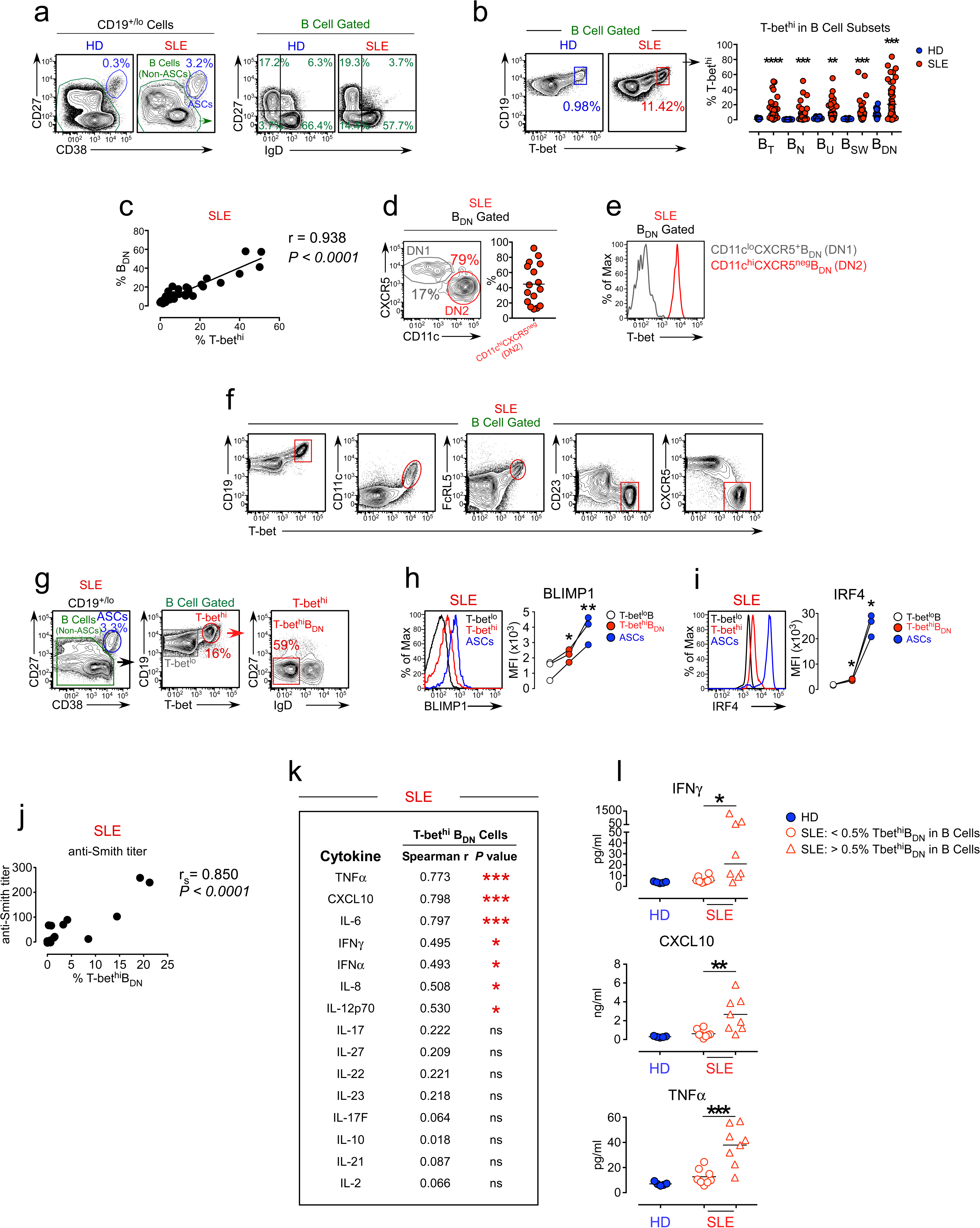
Expansion of the T-bet^hi^ DN2 subset in SLE patients correlates with systemic inflammatory cytokine levels. (**a-c**) Analysis of T-bet^hi^ B cells in peripheral blood B cell subsets from healthy donor (HD) and SLE patients. Gating strategy to identify ASCs, B cells (non-ASCs) (**a, left**) and B cell subsets (**a, right**), including naïve IgD^+^CD27^neg^ (B_N_), switched memory IgD^neg^CD27^+^ (B_SW_) unswitched memory IgD^+^CD27^+^ (B_U_) and double negative IgD^neg^CD27^neg^ (B_DN_) cells, from the peripheral blood of HD and SLE patients. Frequency of T-bet^hi^ B cells (**b**) within the total B cell (B_T_) compartment and each B cell subset with representative flow plot showing T-bet expression in total SLE B cells. Correlation analysis (**c**) comparing frequencies of B_DN_ cells to T-bet^hi^ B cells in each patient. n= 20 HD (**b**) and 40 SLE patients (**b-c**). (**d-e**) T-bet expression by SLE patient B_DN_ cells. Subdivision of the SLE B_DN_ population (**d**) into CXCR5^+^CD11c^lo^ DN1 and T-bet^hi^ CXCR5^neg^CD11c^hi^ DN2 populations with T-bet expression levels (**e**) in each subset shown as a histogram. Data include representative flow plots from a single patient (**d-e**) and the frequency of DN2 cells (**e**) within the B_DN_ subset of 16 SLE patients. (**f**) Phenotypic characterization of T-bet^hi^ B cells in SLE patients. Expression of CD19, CD11c, FcRL5, CD23 and CXCR5 by T-bet^hi^ B cells from a representative SLE patient. (**g-i**) *Ex vivo* isolated SLE T-bet^hi^ B_DN_ cells resemble pre-ASCs. SLE patient B cells (n=3 independent donors) were subdivided as shown in (**g**) into CD27^hi^CD38^hi^ ASCs, T-bet^hi^ B_DN_ cells and T-bet^lo^ B cells and analyzed by flow cytometry for expression of BLIMP1 (**h**) and IRF4 (**i**). Representative flow plots and mean fluorescence intensity (MFI) expression of BLIMP1 and IRF4 in each population are shown. (**j**) Correlation analysis between frequency of circulating T-bet^hi^ B_DN_ cells and anti-Smith autoAb titers in SLE patients. n= 18 SLE patients. (**k-l**) Correlation (**k**) between plasma cytokine levels and frequency of T-bet^hi^ B_DN_ cells in SLE patient peripheral blood. Plasma concentration of IFNγ, CXCL10 and TNFα (**l**) in HD (blue symbols) and SLE patients (red symbols). SLE samples were subdivided into patients with <0.5% (circles) or >0.5% (triangles) T-bet^hi^ B_DN_ cells within the total B cells. n= 5 HD and 16 SLE patients. Individual human subjects in each analysis are represented by a symbol. Horizontal black lines represent the mean (**d** and **l** (CXCL10, TNFα) or median (**b, l** (IFNγ)) within the group. Statistical analyses were performed using a Student’s t test (**l** (CXCL10 and TNFα)), non-parametric Mann-Whitney test (**b** and **l** (IFNγ)), a one way paired T test (**h-i**), Spearman Correlation test (**j-k**) or Pearson Correlation test (**c**). Correlation *P* and r values listed in the figure. *P* values * ≤0.05, **<0.01, ***<0.001.

### Expansion of T-bet^hi^ DN2 cells correlates with systemic IFNγ levels in SLE patients

Since we previously showed that T-bet expression is induced by IFNγ in B cells (42), we hypothesized that the expansion of the T-bet^hi^ DN2 pre-ASC subset in SLE patients would be associated with IFNγ levels in these patients. To test this possibility, we measured 15 cytokines in plasma from the SLE patients. Consistent with our hypothesis, we observed a significant positive correlation between IFNγ, as well as the IFNγ-induced cytokines CXCL10, IL-6 and TNFα, and the frequency of T-bet^hi^ DN2 cells in these individuals (Fig. 1k-l). These data therefore indicated that the T-bet^hi^ DN2 pre-ASC population is most expanded in SLE patients who also exhibit elevated expression of IFNγ and IFNγ-driven inflammatory cytokines.

### IFNγ-producing Th1 cells promote development of T-bet^hi^ B_DN_ cells

Recent experiments from our lab revealed that mouse B cells that are activated in the presence of IFNγ-producing T cells differentiate into ASCs in an IFNγ and T-bet dependent fashion (43). Since human T-bet^hi^ DN2 pre-ASCs are expanded in SLE patients with higher systemic levels of IFNγ, we predicted that the IFNγ might drive the development of the human T-bet^hi^ DN2 pre-ASC population. To begin testing this prediction, we developed an *in vitro* B cell/T cell mixed lymphocyte reaction (MLR) paired co-culture system (Fig. 2a) containing B_N_ cells (Fig. 2b) purified from the peripheral blood or tonsil of one HD and highly polarized human Th1 and Th2 effectors (44), which were generated *in vitro* using purified naïve peripheral blood T cells isolated from a second unrelated HD. The Th1 cells expressed T-bet (Fig. 2c) and produced IFNγ and IL-8 following restimulation (Fig. 2d) while Th2 cells expressed GATA-3 (Fig. 2c) and produced elevated levels of IL-4, IL-5, and IL-13 (Fig. 2d). Since neither the Th1 nor Th2 cells expressed Bcl6 (not shown) or produced IL-21 following restimulation (Fig. 2e), we added IL-21 to the co-cultures to ensure optimal B_N_ activation (45, 46) and included IL-2 to enhance the survival of the T effectors (47). After 6 days in culture, both B cells and ASCs were detected in both cultures (Fig. 2f). Few of the HD B cells activated with IL-4 producing Th2 cells upregulated T-bet (<3%), while more than half of B cells activated in the presence of IFNγ-producing Th1 cells expressed T-bet (Fig. 2f). Approximately 50% of the T-bet^hi^ B cells present in the Be1 cultures downregulated IgD and these cells were CD27^neg^CD19^hi^CD11c^+^FcRL5^+^CD23^neg^ (Fig. 2g). Therefore, activation of B_N_ cells with Th1 cells and IL-21 + IL-2 resulted in the formation of a T-bet^hi^ B_DN_ population that was phenotypically similar to the SLE patient-derived T-bet^hi^ DN2 cells.

**Figure 2.**
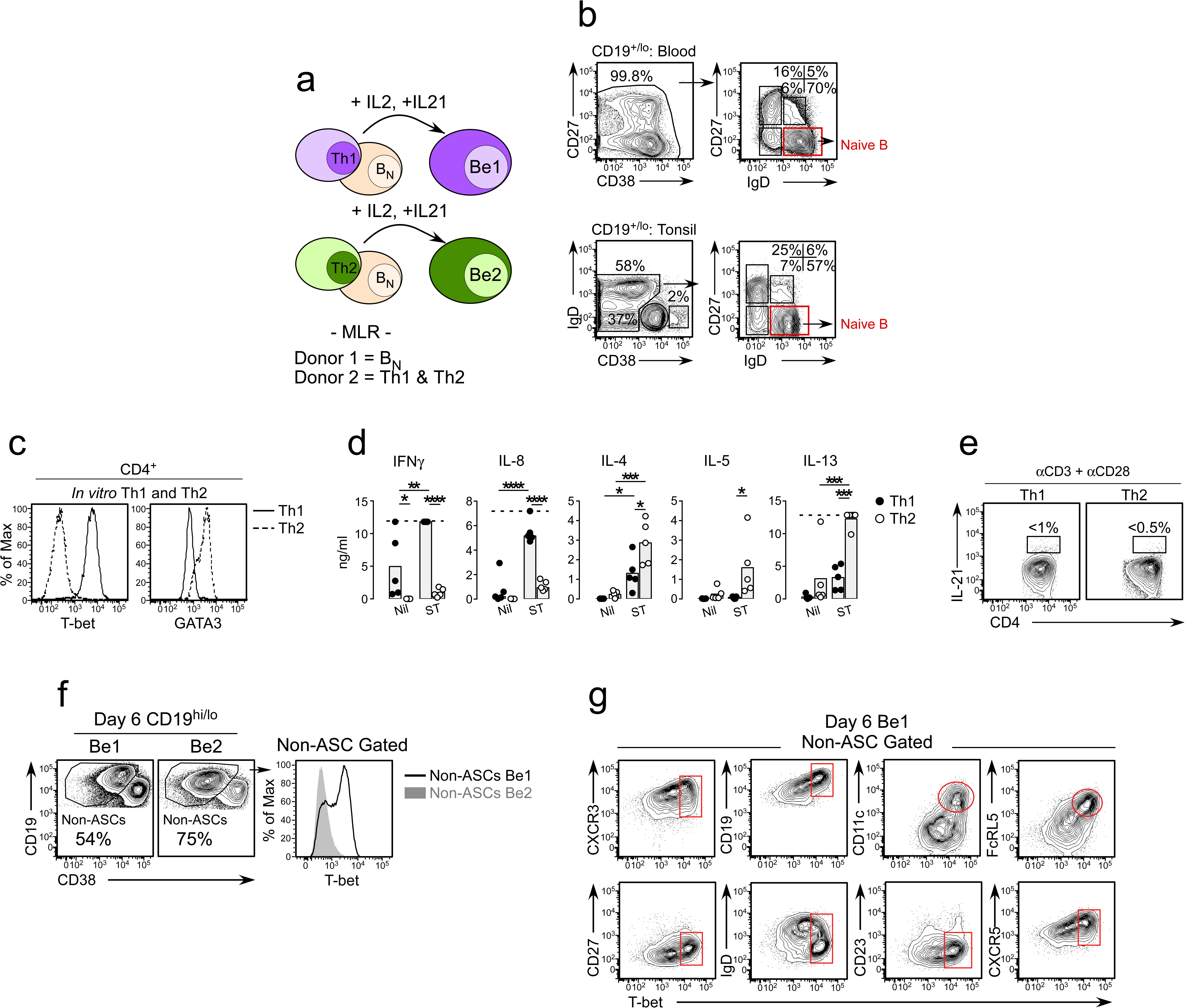
Th1 cells promote the formation of T-bet^hi^ IgD^neg^CD27^neg^ B_DN_ cells from B_N_ precursors. (**a-e**) Description of B/T co-cultures. Cartoon (**a**) describing setup of Be1 (B_N_ + Th1 cells) and Be2 (B_N_ + Th2 cells) co-cultures. Paired Be1 and Be2 co-cultures contain Th1 or Th2 effectors generated from the same HD, B_N_ cells from a second allogeneic HD and exogenous IL-21 and IL-2. Purification strategy (**b**) for B_N_ cells (red gate) from tonsil and blood and characterization of *in vitro* polarized Th1 and Th2 cells (**c-e**). T-bet and GATA-3 expression (**c**) in Th1 (solid line) and Th2 (dotted line) cells restimulated for 6 hr with plate-bound anti-CD3 and anti-CD28. Cytokine levels (**d**) in supernatants from restimulated (ST) or non-restimulated (nil) Th1 (black circles) and Th2 cells (open circles) from 5 independent experiments with the gray bars representing the mean of all experiments. Dotted line indicates maximal measurable levels of the cytokine in the assay. IL-21 production (**e**) by restimulated Th1 and Th2 cells. (**f-g**) Development of T-bet^hi^ B_DN_ cells in B_N_/Th1 co-cultures. Flow cytometric analysis showing T-bet expression (**f**) on gated HD B cells from day 6 Be1 and Be2 co-cultures. Phenotyping (**g**) of day 6 B cell-gated Be1 cells showing T-bet expression in combination with other surface markers. Statistical analyses (**d**) were performed using One-way ANOVA comparing the mean of the non-restimulated to restimulated cells. *P* values *<0.05, **<0.01, ***<0.001, ****<0.0001.

### Differentiation of naïve B cells into ASCs is enhanced in the presence of Th1 cells

Given that the Th1-induced T-bet^hi^ B_DN_ cells were phenotypically related to the previously characterized SLE patient T-bet^hi^ DN2 pre-ASCs, we predicted that the *in vitro* generated T-bet^hi^ B_DN_ subset might represent a pre-ASC population. To test this hypothesis, we first enumerated CD38^hi^CD27^hi^ ASCs in day 6 Be1 and Be2 co-cultures. Although we detected ASCs in both co-cultures (Fig. 3a), we always found more ASCs in the Be1 co-cultures (Fig. 3a), even across multiple independent experiments using B_N_ and T effectors from different HD pairs (Fig. 3b). To address whether the increased ASC formation observed in the Be1 co-cultures was limited to isotype switched or unswitched B cells, we measured the frequency of IgM and IgG-producing (Fig. 3c-d with isotype gating shown in Figure 3-figure supplement 1k) ASCs across multiple paired Be1 and Be2 co-cultures. Again, we found that ASCs, regardless of isotype, were greatly enriched in the Be1 co-cultures (Fig. 3c-d). These data therefore indicated that, while both Be1 and Be2 co-cultures promote ASC formation, Be1 co-culture conditions appear to be highly conducive to ASC development.

**Figure 3.**
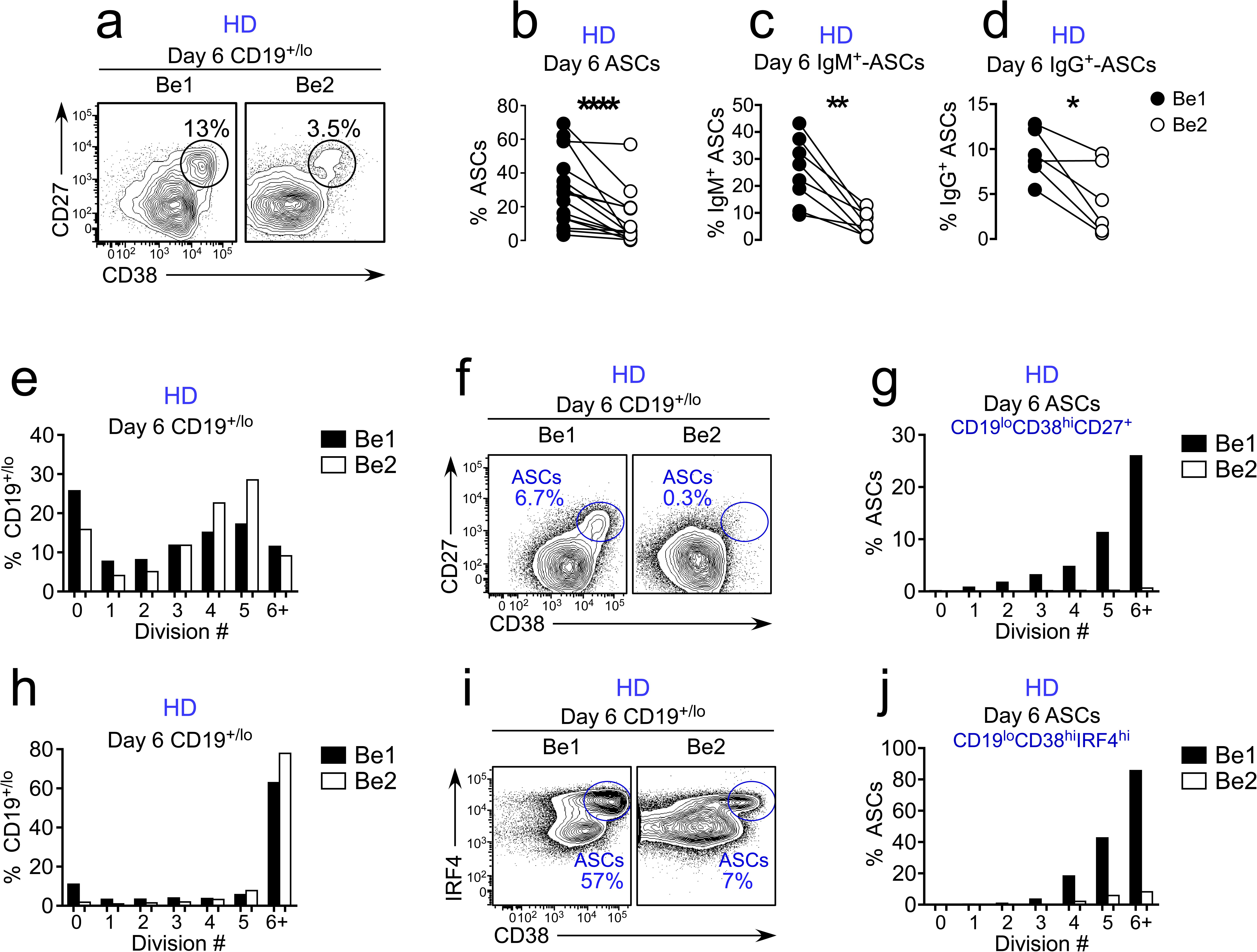
ASC development from B_N_ precursors is enhanced in Th1 containing co-cultures. (**a-d**) ASC development in HD Be1 and Be2 co-cultures showing representative flow plots (**a**) and frequencies (**b**) of CD38^hi^CD27^hi^ ASCs in CD19^+/lo^-gated B lineage cells from day 6 paired Be1 and Be2 co-cultures. Frequencies of IgM^+^ (**c**) or IgG^+^ (**d**) ASCs in each day 6 paired co-culture. Gating strategy to identify IgG^+^ and IgM^+^ ASCs shown in Figure 3-figure supplement 1. (**e-j**) Proliferation analysis of B cells in paired day 6 HD Be1 and Be2 co-cultures. Co-cultures generated with purified CTV-labeled HD B_N_ cells and allogeneic Th1 or Th2 cells + IL-21 and IL-2. B lineage cells gated as CD19^+/lo^ (includes both ASCs and non-ASC B cells). ASCs identified as CD27^hi^CD38^hi^ (**f**) or CD38^hi^IRF4^hi^ (**i**). Data reported as the proportion of total CD19^+/lo^ B lineage cells (**e, h**) in each cell division or the fraction of cells within each cell division that are ASCs (**g, j**). Data from 3 additional independent B_N_/Th1 co-cultures can be found in Figure 3-figure supplement 1. Analyses in **b-d** performed on 15 (**b**), 8 (**c**) or 6 (**d**) independent paired Be1 and Be2 co-cultures. Analysis in **e-j** are from two independent paired co-cultures and are representative of >5 independent experiments (See Figure 3-figure supplement 1). Statistical analyses were performed using a non-parametric Wilcoxon paired t test (**b**) or paired Student’s t test (**c-d**). *P* values *<0.05, **<0.01, ****<0.0001.

To determine whether the increased ASC formation in the Be1 cultures was due to increased proliferation, we set up paired Be1 and Be2 co-cultures with Cell Trace Violet (CTV)-labeled B_N_ cells and monitored proliferation and ASC formation in the cultures. As shown in Figure 3e, the B cells proliferated in both cultures, with a similar frequency of B cells represented in each cell division. However, despite equivalent rates of proliferation, the frequency of CD27^hi^CD38^hi^ ASCs was ∼20-fold higher in the Be1 co-culture compared to the paired Be2 co-culture (Fig. 3f). In fact, 40% of the B cells that had divided at least 5 times in the Be1 cultures were ASCs while <2% of the B cells that had divided ≥5 times were ASCs in the Be2 cultures (Fig. 3g). The same analysis was performed in an additional four independent paired Be1 and Be2 co-cultures (Fig. 3h-j and Figure 3-figure supplement 1a-i) and, while replicative response in each of the independent allo co-cultures was unique, we always observed equivalent proliferative rates between the Be1 and Be2 cells in the co-cultures and enhanced ASC formation in the Be1 cultures (mean 18.35-fold and median 11.2-fold increase in percentage of ASCs in Be1 co-cultures). Importantly, the ASCs that were found in each co-culture exhibited the same proliferative history with 89% of the ASCs present in Be1 cultures having undergone ≥5 divisions and 88% of the ASCs present in Be2 cultures having divided ≥5 times (Figure 3-figure supplement 1j). Therefore, we conclude that the increased ASC formation in Be1 cultures compared to the Be2 cultures is not due to intrinsic differences in the proliferative rates of the cells in each culture but rather that a higher proportion of the Be1 cells at each cell division make the commitment to the ASC lineage.

### T-bet^hi^ B_DN_ cells induced with Th1 cells and IL-21 are pre-ASCs

Given the phenotypic similarities between the *in vitro* induced T-bet^hi^ B_DN_ cells and T-bet^hi^ DN2 cells from SLE patients and the fact that the *in vitro* cultures containing T-bet^hi^ B_DN_ cells also efficiently formed ASCs, we predicted that the Tbet^hi^ B_DN_ cells found in the Th1/B_N_ co-cultures were likely to be pre-ASCs. To test this, we first asked whether the *in vitro* generated Th1-induced T-bet^hi^ B_DN_ cells were transcriptionally related to T-bet^hi^ DN2 pre-ASCs from SLE patients. We therefore sort-purified IgD^neg^CD27^neg^ B_DN_ cells (Fig. 4a) from three independent paired day 6 Be1 and Be2 co-cultures and performed RNA-seq analysis. We identified 427 differentially expressed genes (DEGs) between the B_DN_ cells from the Be1 and Be2 co-cultures (Fig. 4b, Supplementary File 1). Consistent with our data showing that T-bet was selectively upregulated in the B cells from Be1 co-cultures, we observed significantly higher levels of *TBX21* mRNA in the *in vitro* induced B_DN_ Be1 cells compared to B_DN_ Be2 cells (Fig. 4c). Next, we used Gene Set Enrichment Analysis (GSEA) to compare the transcriptomes of the *in vitro* generated B_DN_ cells isolated from the Be1 and Be2 cultures with the transcriptome of the T-bet^hi^ DN2 population isolated from SLE patients ((19), Supplementary File 2). Consistent with our phenotyping data, the transcriptome of the T-bet expressing B_DN_ Be1 cell subset was highly enriched relative to the B_DN_ Be2 cells for genes that are differentially upregulated in the SLE-derived T-bet^hi^ DN2 subset (Fig. 4d). Similarly, the transcriptome of the *in vitro* generated T-bet^hi^ B_DN_ Be1 cells was enriched in genes that are upregulated in the CD11c^+^ T-bet expressing ABCs (48) isolated from aged mice (Fig. 4e-f).

**Figure 4.**
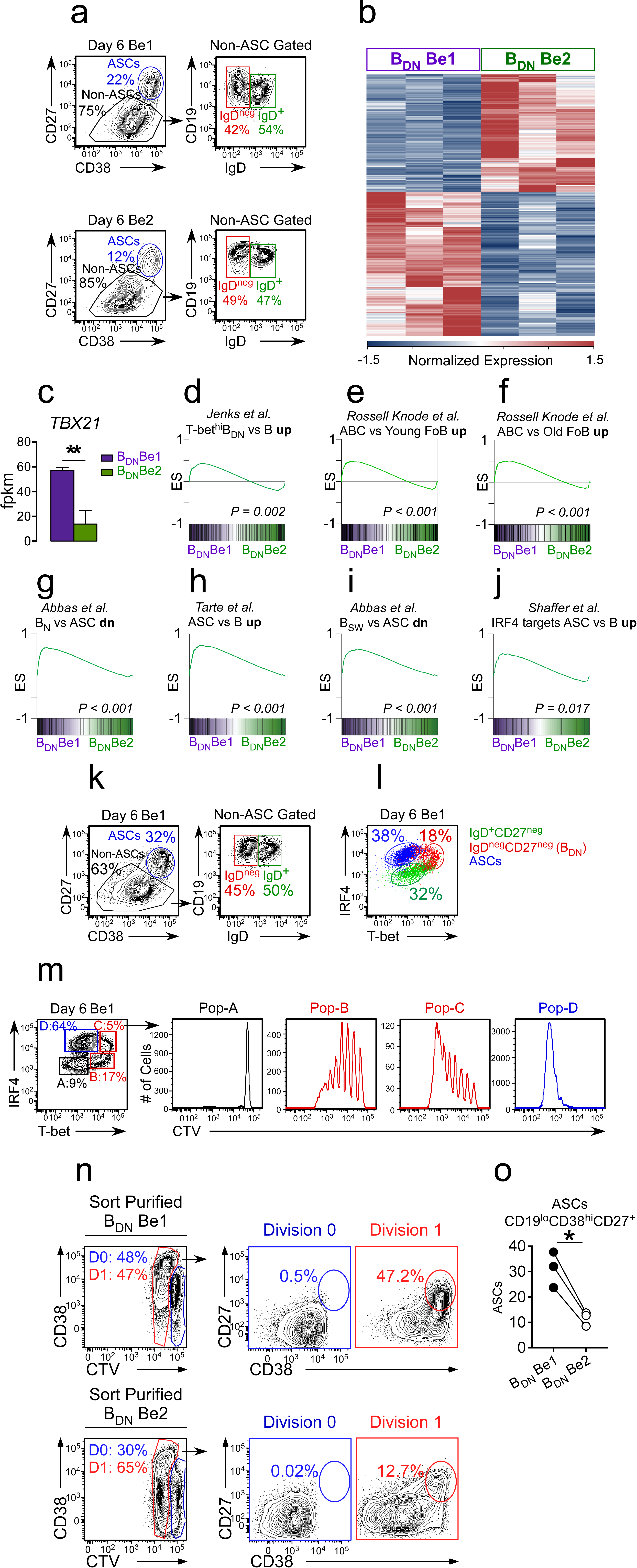
Th1-induced T-bet^hi^ B_DN_ cells are pre-ASCs. (**a-j**) Transcriptome analysis of *in vitro* generated IgD^neg^CD27^neg^ T-bet^hi^ B_DN_ cells. RNA-seq analysis performed on IgD^neg^CD27^neg^ B_DN_ cells (gating in panel **a**) that were sort-purified from day 6 HD Be1 and Be2 co-cultures. (n=3 samples/subset derived from 3 independent paired co-culture experiments). Heat map (**b**), showing 427 differentially expressed genes (DEGs) based on FDR < 0.05. T-bet mRNA expression levels (**c**) in IgD^neg^CD27^neg^ B_DN_ cells from day 6 Be1 and Be2 co-cultures. Gene Set Enrichment Analysis (GSEA, panels **d-j**) comparing transcriptome profile of *in vitro* generated B_DN_ cells from Be1 and Be2 co-cultures with published DEGs identified in different B cell subsets. Data are reported as Enrichment Score (ES) plotted against the ranked B_DN_ Be1 and Be2 gene list (n = 11598). DEG lists used for GSEA include: DEGs that are upregulated in sort-purified SLE patient-derived T-bet^hi^ B_DN_ cells (CD19^hi^IgD^neg^CD27^neg^CXCR5^neg^IgG^+^) compared to other SLE patient-derived mature B cell subsets (**d,** (19)); DEGs that are upregulated in CD11c^hi^ ABCs isolated from aged mice (48) compared to follicular B cells isolated from young (**e**) or aged (**f**) mice; DEGs that are upregulated in human plasma cells (ASCs) relative to: B_N_ cells (**g**, (49)), total B cells (**h**, (50)) or B_SW_ cells (**i**, (49)); and IRF4 target genes (**j**, (51)) that are upregulated in ASCs relative to B cells. Nominal *P* value for GSEA is shown. (**k-m**) IgD^neg^CD27^neg^ T-bet^hi^ B_DN_ cells express intermediate levels of IRF4 and are actively proliferating. Gating strategy (**k**) to identify CD27^hi^CD38^hi^ ASCs, IgD^+^CD27^neg^ B cells and IgD^neg^CD27^neg^ B_DN_ cells in day 6 Be1 co-cultures generated from unlabeled (**k-l**) or CTV-labeled (**m**) HD B_N_ cells. Expression of T-bet and IRF4 (**l-m**) by ASCs (blue), IgD^+^CD27^neg^ B cells (green) and IgD^neg^CD27^neg^ B_DN_ cells (red) from day 6 Be1 co-cultures. Proliferation profile of T-bet^lo^IRF4^neg^ (Pop-A), T-bet^hi^IRF4^lo/int^ (Pop-B), T-bet^hi^IRF4^int/hi^ (Pop-C) and T-bet^lo^IRF4^hi^ (Pop-D) subsets shown. (**n-o**) IgD^neg^CD27^neg^ B_DN_ Be1 cells rapidly differentiate into ASCs. IgD^neg^CD27^neg^ B_DN_ cells from day 6 HD Be1 and Be2 cultures were sort-purified, CTV labeled and incubated overnight (18 hrs) in conditioned medium. Enumeration of ASCs (CD19^lo^CD38^hi^CD27^+^) in the undivided cells (Division 0 (D0)) and the cells that divided one time (Division 1 (D1)). Representative flow plots (**n**) showing the frequency of cells in D0 or D1 in each culture and the frequency of CD19^lo^CD38^hi^CD27^+^ ASCs present in the D0 or D1 fraction. Panel (**o**) reports frequency of ASCs within the cultures from 3 independent experiments. Statistical analysis performed with unpaired (**c**) or paired (**o**) Students t test. Nominal *P* values (**d-j**) for GSEA are shown. *P* values *<0.05, **<0.01. See Supplementary File 1 for complete B_DN_ Be1 and Be2 RNA-seq data set and Supplementary File 2 for SLE patient-derived T-bet^hi^ B_DN_ DEG list.

Next, we used GSEA to compare the transcriptional profile of the *in vitro* generated B_DN_ cells isolated from the Be1 and Be2 cultures with curated ASC transcriptome datasets (49, 50). Interestingly, the transcriptome of the *in vitro*-induced B_DN_ Be1 population was significantly enriched in expression of genes that are upregulated in ASCs compared to B_N_ cells (Fig. 4g), mature B cells (Fig. 4h) and B_SW_ cells (Fig. 4i). In addition, genes that are direct targets of IRF4 and upregulated in ASCs (51) were significantly enriched in the *in vitro* generated Be1 B_DN_ cells relative to the Be2 B_DN_ cells (Fig. 4j). Consistent with this finding, we observed that the Be1 T-bet^hi^ B_DN_ cells express intermediate levels of IRF4, when compared to the CD27^hi^CD38^hi^ ASCs present and the IgD^+^CD27^neg^ B cells that are present in the Be1 cultures (Fig. 4k-l). Furthermore, when we performed the same experiment using CTV-labeled B_N_ cells, it was clear that the IRF4 expression levels were tied to the proliferative history of the T-bet^hi^ B_DN_ cells (Fig. 4m) and that the Tbet^hi^IRF4^int^ B_DN_ subset was a potential precursor of the T-bet^lo^IRF4^hi^ ASCs.

To confirm that the Be1 T-bet^hi^ B_DN_ cells are functional pre-ASCs, we sort-purified the IgD^neg^CD27^neg^ B_DN_ cells from both Be1 and Be2 co-cultures, labeled the sorted subsets with CTV, incubated the cells for 18 hrs in conditioned media and finally enumerated CD27^hi^CD38^hi^ ASCs in the cultures. As expected, the sorted Be1 and Be2 B_DN_ cells were activated with 47-65% of the cells undergoing one cell division within 18 hrs (Fig. 4n). CD27^hi^CD38^hi^ ASCs were only detected in proliferating cells (Fig. 4n), indicating that the sorted B_DN_ cells include pre-ASCs that are poised to differentiate within one round of replication. Importantly, while both Be1 and Be2 B_DN_ cells gave rise to ASCs, ASC development was significantly enhanced in cultures containing the sorted T-bet expressing Be1 B_DN_ cells (Fig. 4o). Thus, activation of B_N_ cells with Th1 cells and IL-21 + IL-2 gives rise to a population of T-bet^hi^ B_DN_ cells that are similar at a phenotypic, molecular and functional level to the T-bet^hi^ DN2 pre-ASCs that are expanded in SLE patients (19).

### Identification of predicted regulators of the T-bet^hi^ B_DN_ pre-ASC population

Given the similarities between the *in vitro* generated Th1-induced T-bet^hi^ B_DN_ subset and the T-bet^hi^ DN2 pre-ASC population that is expanded in some SLE patients (19), we hypothesized that we could use the transcriptome data from our *in vitro* generated pre-ASCs to could be used to predict upstream molecular signals that might give rise to this these cells in SLE patients. We therefore analyzed the RNA-seq data from the Be1 and Be2 B_DN_ cells using Ingenuity Pathway Analysis (IPA) to identify predicted upstream regulators that direct B_N_ cells to develop into T-bet^hi^ B_DN_ pre-ASCs. Not unexpectedly, predicted upstream regulators of the *in vitro* generated Be1 B_DN_ cells included type 1 and type 2 IFNs and the IFN-induced transcription factor STAT1 (Fig. 5a). Interestingly, and despite the fact that exogenous IL-2 + IL-21 and matched allogeneic T cells were included in both Be1 and Be2 cultures, Ag receptor signals, IL-2 and the IL-21 activated transcription factor STAT-3 were predicted to be upstream activators of the Be1 B_DN_ cells but not the Be2 B_DN_ cells (Fig. 5a). In addition, both TLR7 and TLR9 were predicted as upstream regulators of the T-bet^hi^ B_DN_ Be1 cells (Fig. 5a). This was surprising, given that we did not add exogenous TLR ligands to the co-cultures, however, endogenous TLR ligands are known to be released by dying cells *in vitro* (52). Collectively, these data suggested that the T-bet^hi^ B_DN_ cells might be hyperresponsive to IL-2, IL-21 and/or TLR ligands, similar to what was reported for T-bet^hi^ DN2 cells from SLE patients (19).

**Figure 5.**
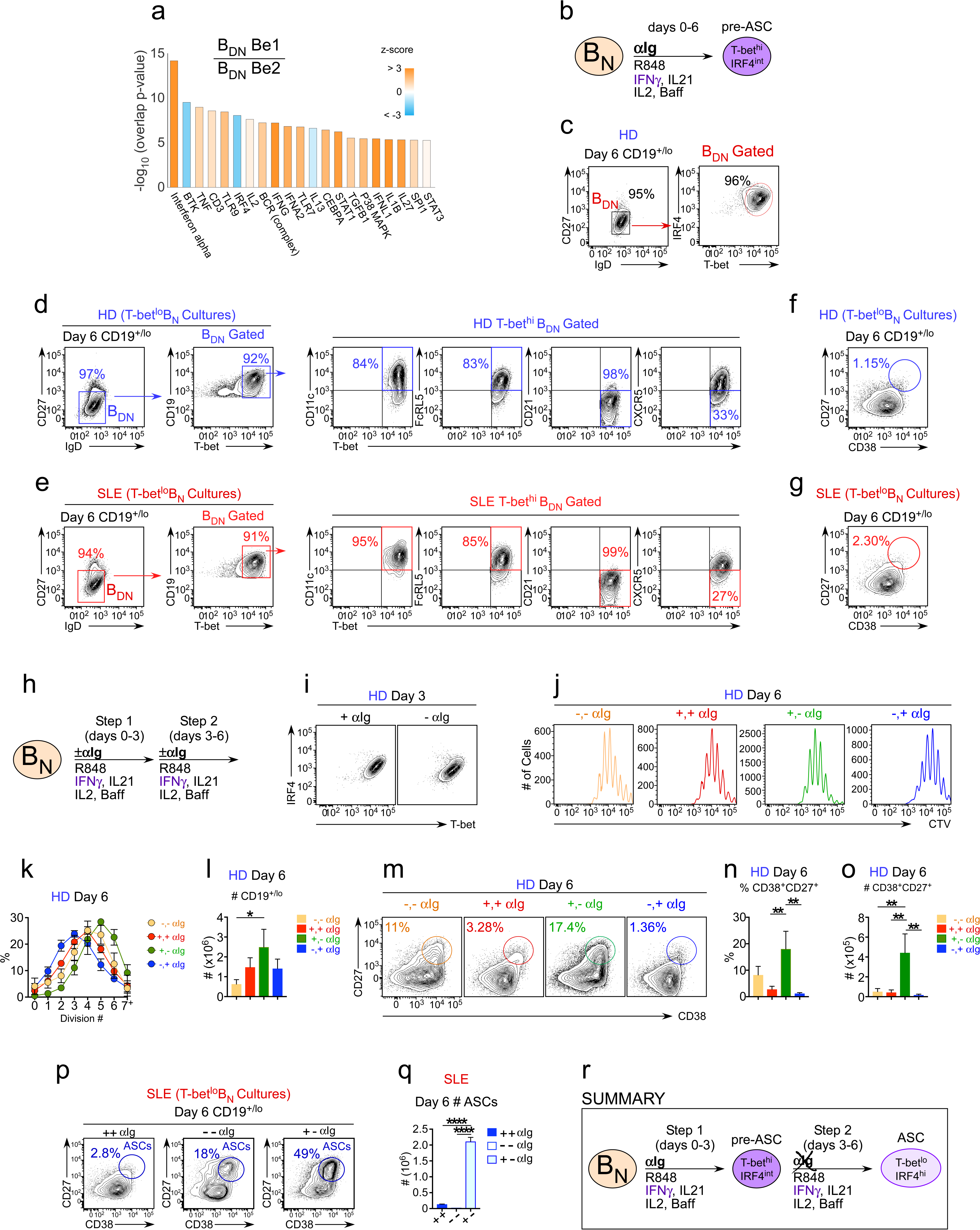
Early but transient BCR signals promote ASC differentiation from T-bet^hi^ B_DN_ pre-ASCs. (**a**) Ingenuity Pathway Analysis (IPA) based on 427 DEG with FDR < .05 (allowing both direct and indirect interactions) to identify predicted upstream regulators of the HD B_DN_ Be1 transcriptome. The predicted activation state (z-score of B_DN_ Be1 over B_DN_ Be2) of each regulator/signaling pathway is shown as bar color (orange, activated; blue, inhibited) with predicted upstream regulators sorted in order of significance (overlap *P* value). Regulators are shown that have an overlap *P*-value < 0.00001. (**b-g**) IPA-identified stimuli induce development of T-bet^hi^IRF4^int^ B_DN_ pre-ASC-like cells from HD and SLE B_N_ cells. Cartoon (**b**) depicting *in vitro* stimulation conditions to activate purified B_N_ cells from HD (**c-d, f**) and SLE patients (**e,g**) with cytokines (IL-2, Baff, IL-21, IFNγ), anti-Ig and R848 for 6 days. Phenotypic characterization of day 6 activated cells showing IRF4 and T-bet expression (**c**) on HD IgD^neg^CD27^neg^ B_DN_ cells. Phenotypic analysis of T-bet^hi^ B_DN_ cells in day 6 cultures containing B_N_ cells from HD (**d**) or SLE (**e**) patients. Enumeration of ASCs in day 6 cultures containing B_N_ cells from HD (**f**) or SLE (**g**) patients. (**h-o**) ASC generation from HD B_DN_ cells requires early but transient BCR activation. CTV-labeled HD B_N_ cells were activated for 3 days with R848, cytokines (IFNγ, IL-2, IL-21, Baff) ± anti-Ig (Step 1) and then washed and recultured for an additional 3 days with the same stimuli ± anti-Ig (Step 2) (**h**). Cells were analyzed by flow cytometry on days 3 (**i**) and 6 (**j-o**). T-bet and IRF4 expression (**i**) on day 3 by R848 and cytokine cocktail activated B cells exposed (+) or not (-) to anti-Ig over the first 3 days. CTV dilution and cell division (**j-k**), cell recovery (**l**), ASC frequencies (**m-n**) and ASC recovery (**o**) on day 6 in cultures that were not exposed to anti-Ig during steps 1 and 2 (-,-); were exposed to anti-Ig throughout steps 1 and 2 (+,+); were exposed to anti-Ig only in step 1 (+,-); or were exposed to anti-Ig only in step 2 (-,+). Representative flow panels showing CTV dilution (**j**) with quantitation showing % of cells in each cell division (**k**) and cell recovery (**l**). Representative flow panels showing % ASCs (**m**) with quantitation showing % ASCs (**n**) and ASC recovery (**o**). (**p-q**) ASC generation from SLE B_N_ cells requires early but transient BCR activation. CTV-labeled SLE T-bet^lo^ B_N_ cells were sort-purified and cultured for 6 days as described in (**h**). Data are shown as the frequency of CD27^hi^CD38^hi^ ASCs (**p**) and ASC recovery (**q**) in cultures containing anti-Ig for the indicated time ((+,+), (-,-) or (+,-)). Summary of data (**r**) showing that ASC development from T-bet^hi^IRF4^int^ B_DN_ population requires removal of anti-Ig from the cultures between days 3-6. RNA-seq IPA analysis was performed on n=3 samples/subset derived from 3 independent paired co-culture experiments. Data in **c-o** are representative of ≥3 experiments. The percentage of cells in each division, the frequency of ASCs and cell recovery (total and ASCs) are shown as the mean ± SD of cultures containing purified B_N_ cells from 3 independent donors. Data shown in (**p-q**) are from a single SLE individual and are representative of 2 independent experiments. Statistical analyses (**l, n, o, q**) were performed using one-way ANOVA with Tukey’s multiple comparison test. *P* values *<0.05, **<0.01, ***<0.001, ****<0.0001.

### Transient BCR stimulation promotes ASC development from T-bet^hi^ B_DN_ pre-ASCs

Using the predictions from the IPA analysis of the Be1 T-bet^hi^ B_DN_ cells we next asked whether we could induce the formation of the T-bet^hi^ B_DN_ pre-ASC population using fully defined stimuli. We therefore activated HD B_N_ cells for 6 days with anti-Ig, cytokines (IFNγ, IL-2, IL-21 and Baff) and the TLR7/8 ligand, R848 (Fig. 5b). Greater than 95% of the B cells activated with these defined stimuli were IgD^neg^CD27^neg^ T-bet^hi^IRF4^int^ (Fig. 5c). In addition, these cells expressed CD11c and FcRL5 but not CD21 and had begun downregulating CXCR5 (Fig. 5d) and were thus phenotypically similar to the SLE patient T-bet^hi^ DN2 cells (19). Importantly, we obtained similar results when we stimulated sort-purified T-bet^lo^ B_N_ cells from SLE patients with the same activation cocktail (Fig. 5e), suggesting that these defined stimuli are sufficient to activate T-bet^lo^ B_N_ cells, isolated from either HD or SLE patients, to develop into a population of T-bet^hi^ B_DN_ cells that are phenotypically similar to the SLE patient-derived T-bet^hi^ DN2 population.

Despite the fact that >95% of the HD or SLE B_N_ cells activated with these defined stimuli developed into what phenotypically appeared to be T-bet^hi^ B_DN_ pre-ASCs (Fig. 5d-e), no CD38^hi^CD27^hi^ ASCs were detected in either culture (Fig. 5f-g). This suggested to us that our cultures were either missing a stimulus that is required for the differentiation of the pre-ASCs into ASCs or that one of the factors in our defined stimulation cocktail prevents differentiation of T-bet^hi^ B_DN_ cells into ASCs. Since our original Th1/B_N_ co-cultures did not contain anti-Ig, we examined whether removing anti-Ig or only providing it transiently would impact pre-ASC and ASC formation. We therefore stimulated CTV-labeled B_N_ cells for six days with the complete activation cocktail (+,+) or removed the anti-Ig from the activation cocktail for the first three days (-,+), last three days (+,-), or throughout the entire culture period (-,-) (Fig. 5h). T-bet^hi^IRF4^int^ cells were easily detected by day 3 in the cultures that lacked anti-Ig for the first 3 days (Fig. 5i), indicating that early BCR signals are not obligate for the development of the pre-ASC population. However, the proliferative response of the B cells (Fig. 5j-k), cell recovery (Fig. 5l) and ASC development (Fig. 5m-n) was highly dependent on early but transient exposure to anti-Ig. Specifically, ASCs were detected when anti-Ig was present during the first three days of culture but were greatly reduced when anti-Ig was added late to the cultures or included for all 6 days (Fig. 5m-n). Moreover, early (day 0-3) but transient stimulation with anti-Ig resulted in more proliferation (Fig. 5j-k), better cell recovery (Fig. 5l) and maximal recovery of ASCs on day 6 (Fig. 5o) compared to all other conditions. To confirm that B_N_ cells from SLE patients behaved similarly, we activated sort purified SLE T-bet^lo^ B_N_ cells with the same stimulation cocktail without any anti-Ig or included anti-Ig for the first 3 days of the culture or throughout the whole culture period. Again, ASC recovery was optimal when anti-Ig was included transiently during the first three days of the culture (Fig. 5p-q). Taken together, the data indicated that early and transient BCR ligation enhanced cell recovery, proliferation and ASC formation when B_N_ cells from either HD or SLE patients were activated with R848, IFNγ, IL-21, IL-2 and Baff, while sustained BCR stimulation suppressed ASC development (Fig. 5q).

### ASC development from T-bet^hi^IRF4^int^ pre-ASCs is regulated by IFNγ, TLR7 and IL-21

Now that we had identified a set of defined stimulation conditions that induced the formation of T-bet^hi^ B_DN_ cells and ASCs, we next asked which signals were critical for the formation of the T-bet^hi^ B_DN_ subset and the development and maximal recovery of ASCs in the cultures. We therefore set up “all minus one cultures” by activating CTV-labeled B_N_ cells for six days – three days in the presence of anti-Ig and three days without anti-Ig – while excluding one stimulus for all six days of the culture (Fig. 6a). As expected, when HD B_N_ cells were activated for three days in the presence of anti-Ig and all cytokines + R848, >90% of the cells upregulated T-bet and IRF4 (Fig. 6b). Similar results were observed when the anti-Ig stimulated B_N_ cells were activated for three days without R848, IL-21, BAFF or IL-2 (Fig. 6b). Thus, despite earlier studies using mouse B cells that showed that T-bet expression can be induced by TLR and IL-21 signals (53), these data show that, at least under this set of stimulation conditions, TLR and IL-21 are not obligate for upregulation of T-bet or IRF4. By contrast, when the cells were activated without IFNγ, more than 80% of the cells were T-bet^neg/lo^. While this wasn’t particularly surprising, given that T-bet is an IFNγ-induced transcription factor (14), the cells also failed to upregulate IRF4 (Fig. 6b). Thus, IFNγ signals are required for the establishment of the T-bet^hi^IRF4^int^ pre-ASC population.

**Figure 6.**
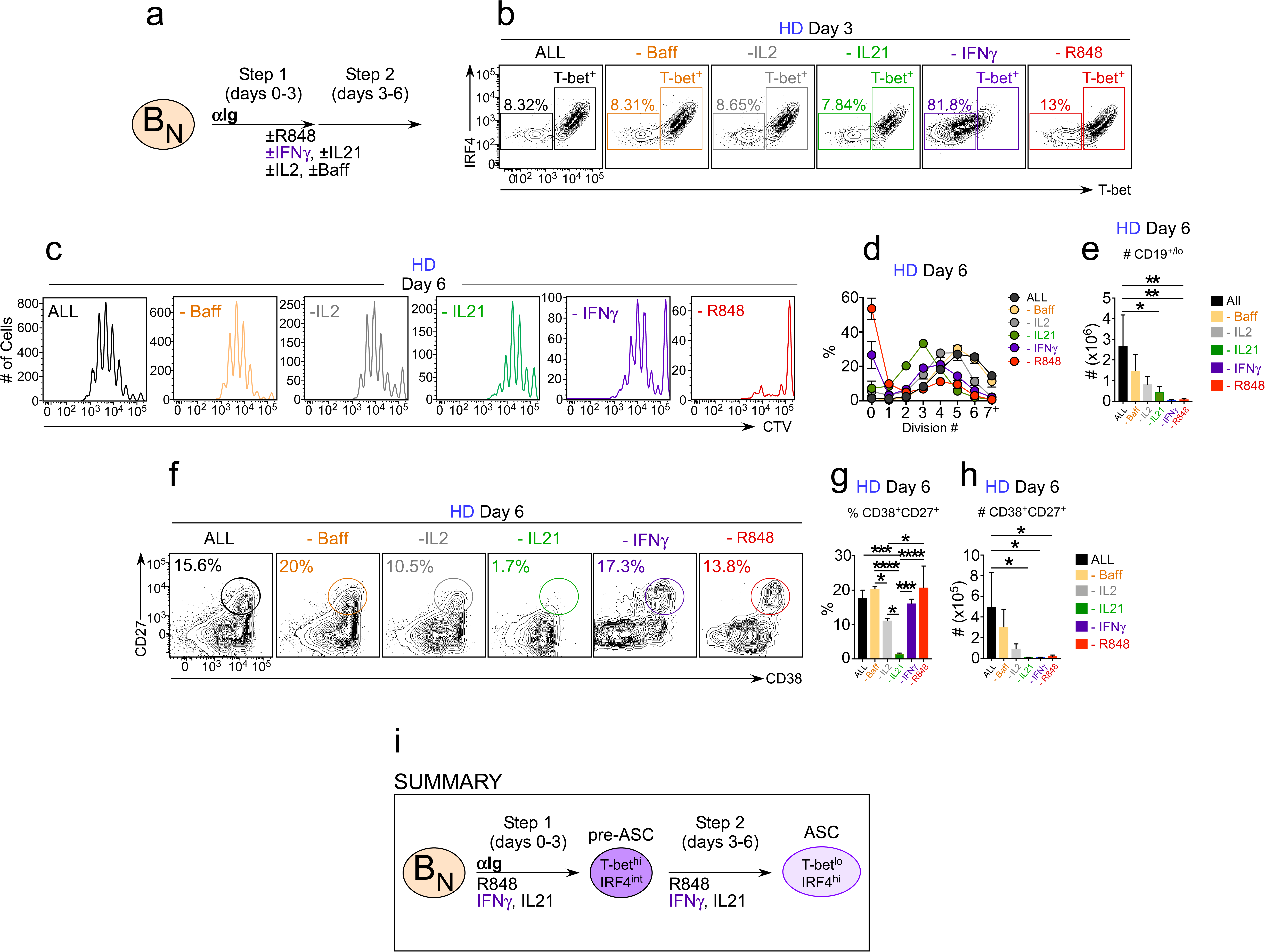
Development of T-bet^hi^IRF4^int^ B_DN_ pre-ASCs and ASCs is controlled by IFNγ, R848, IL-2 and IL-21. CTV-labeled HD B_N_ cells were activated with anti-Ig + cytokine cocktail (IFNγ, IL-2, IL-21, Baff) and R848 for 3 days (Step 1) and then cultured for an additional 3 days (Step 2) with cytokine cocktail and R848 (“ALL” condition). Alternatively, individual stimuli (as indicated) were excluded from the cultures for all 6 days (**a**). Cells were analyzed by flow cytometry on days 3 (**b**) and 6 (**c-h**). T-bet and IRF4 expression (**b**) on day 3 in “ALL” cultures or cultures lacking the indicated stimulus. Representative flow panels showing CTV dilution (**c**) with quantitation showing % of cells in each cell division (**d**) and cell recovery (**e**). Representative flow panels showing % ASCs (**f**) with quantitation showing % ASCs (**g**) and ASC recovery (**h**). Summary of data (**i**) showing that development of the T-bet^hi^IRF4^int^ B_DN_ pre-ASC population requires IFNγ and that recovery of ASCs in the cultures on day 6 is dependent on IFNγ, IL-2, IL-21 and R848. Plots depicting CTV dilution, T-bet^hi^IRF4^int^ pre-ASCs and CD27^hi^CD38^hi^ ASCs in each culture are representative of ≥3 experiments. The percentage of cells in each division, the frequency of ASCs and cell recovery (total and ASCs) are shown as the mean ± SD of cultures containing purified B_N_ cells from 3 independent healthy donors. All statistical analyses were performed using one-way ANOVA with Tukey’s multiple comparison test. *P* values *<0.05, **<0.01, ***<0.001, ****<0.0001.

In agreement with our earlier experiments, we observed maximal cell proliferation (Fig. 6c-d), cell recovery (Fig. 6e) and ASC development (Fig. 6f-g) when B cells were activated with anti-Ig, R848 and cytokines for three days and then incubated an additional three days with the same stimuli minus anti-Ig. Despite the known role for BAFF in mature B cell survival (54), eliminating BAFF from the cultures had only a modest impact on any of the measured parameters (Fig. 6c-g). Consistent with prior reports showing that ASC development from B_N_ cells is dependent on IL-21 (45, 46), few ASCs, whether measured by frequency (Fig. 6f-g) or number (Fig. 6h), were present in the cultures lacking IL-21. Although similar frequencies of ASCs were found in cultures that lacked R848, IFNγ or IL-2 (Fig. 6f-g), proliferation (Fig. 6c-d) and cell recovery (Fig. 6e), including recovery of ASCs on day 6 (Fig. 6h), were significantly impaired. In fact, total ASC recovery in cultures lacking R848 or IFNγ was as low as that observed in the cultures lacking IL-21 (Fig. 6h). These data therefore indicated that BAFF is dispensable for the formation and recovery of pre-ASCs and ASCs. IL-2, while not absolutely essential, contributes significantly to ASC recovery. Finally, IFNγ, R848 and IL-21 play critical but distinct roles in the formation of T-bet^hi^IRF4^int^ pre-ASC population and in the development and recovery of ASCs (Fig. 6i).

### Temporal control of ASC development from T-bet^hi^IRF4^int^ pre-ASCS by IFNγ, TLR7 ligand and IL-21

Our data indicated IFNγ was required for the development of the T-bet^hi^IRF4^int^ pre-ASCs and suggested that ASC formation from this pre-ASC population was promoted by IL-21 and repressed by sustained BCR signaling. Moreover, the data showed that IFNγ and TLR7/8 signals were critically important for ASC recovery. Given these results, we postulated that IFNγ signals would be more important early after activation (Days 0-3, priming phase) while TLR7/8 and IL-21 signals would be more critical later in the culture period (Days 4-6, expansion/differentiation phase). To test this hypothesis, we measured proliferation, cell recovery and the frequency and number of ASCs present in cultures containing CTV-labeled B_N_ cells that were activated for three days in the presence of anti-Ig and three days without anti-Ig – while adding IFNγ (Fig. 7a-h), R848 (Fig. 7i-p) or IL-21 (Fig. 7q-x) during the priming phase (+,-), during the expansion/differentiation phase (-,+) or throughout (+,+) the culture period.

**Figure 7.**
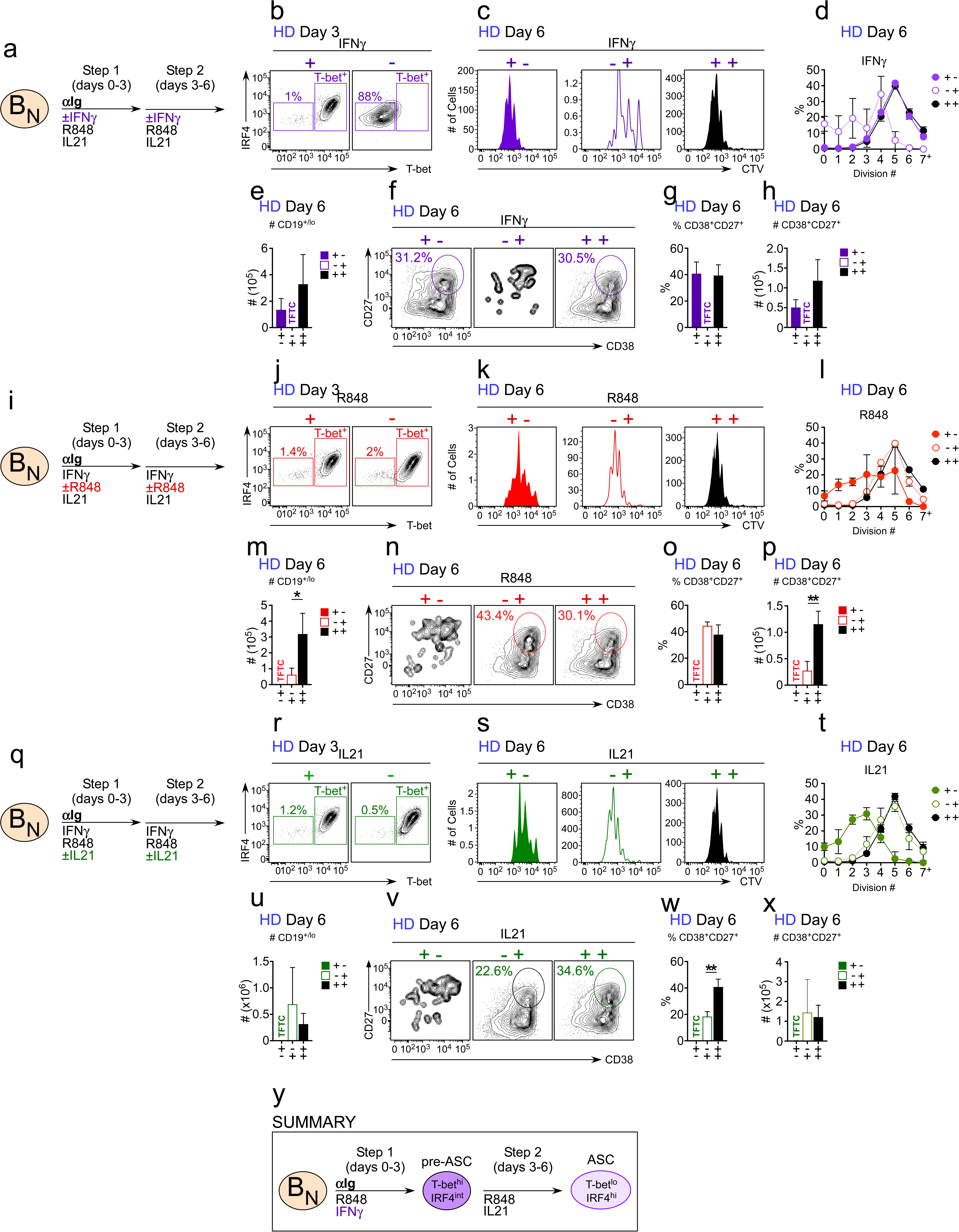
Temporally distinct regulation of T-bet^hi^IRF4^int^ pre-ASC and ASC development by IFNγ, TLR7 ligand and IL21. CTV-labeled HD B_N_ cells were activated for 3 days with anti-Ig, R848, IL-21 and IFNγ (Step 1), washed and then re-cultured for 3 days with R848, IFNγ, and IL-21 (Step 2, +,+ condition). Alternatively, individual stimuli were included in Step 1 only (+,-condition) or in Step 2 only (-,+ condition). (**a-h**) Cells from cultures containing IFNγ in Step 1, Step 2 or both steps (**a**) were analyzed by flow cytometry on days 3 (**b**) and 6 (**c-h**). T-bet and IRF4 expression (**b**) on day 3. CTV dilution and cell division (**c-d**), cell recovery (**e**), ASC frequencies (**f-g**) and ASC recovery (**h**) by day 6 B cells. (**i-p**) Cells from cultures containing R848 in Step 1, Step 2 or both steps (**i**) were analyzed by flow cytometry on days 3 (**j**) and 6 (**k-p**). T-bet and IRF4 expression (**j**) on day 3. CTV dilution and cell division (**k-l**), cell recovery (**m**), ASC frequencies (**n-o**) and ASC recovery (**p**) by day 6 B cells. (**q-x**) Cells from cultures containing IL-21 in Step 1, Step 2 or both steps (**q**) were analyzed by flow cytometry on days 3 (**r**) and 6 (**s-x**). T-bet and IRF4 expression (**r**) on day 3. CTV dilution and cell division (**s-t**), cell recovery (**u**), ASC frequencies (**v-w**) and ASC recovery (**x**) by day 6 B cells. (**y**) Summary of data showing that ASC development and recovery from T-bet^hi^IRF4^int^ B_DN_ pre-ASCs requires early “priming” signals IFNγ and R848 and late proliferation/differentiation signals from R848 and IL-21. Flow cytometry plots, depicting CTV dilution, T-bet^hi^IRF4^int^ pre-ASCs and CD27^hi^CD38^hi^ ASCs in each culture, are representative of ≥3 experiments. The percentage of cells in each division, the frequency of ASCs and cell recovery (total and ASCs) are shown as the mean ± SD of cultures containing purified B_N_ cells from 3 independent healthy donors. All statistical analyses were performed using Student’s T test. *P* values *<0.05, **<0.01, ***<0.001, ****<0.0001. TFTC = Too few to count.

As expected, eliminating IFNγ from the cultures during the first 3 days (Fig. 7a) prevented formation of the T-bet^hi^IRF4^int^ pre-ASC population (Fig. 7b). In addition, B_N_ cells that did not receive an IFNγ signal during the priming phase proliferated less over the 6 day culture period (Fig. 7c-d), resulting in minimal cell recovery on day 6 (Fig. 7e). By contrast, adding IFNγ in the priming phase was sufficient to induce formation of the T-bet^hi^IRF4^int^ pre-ASC population (Fig. 7b) and to promote proliferation and cell recovery measured on day 6 (Fig. 7c-e). Moreover, addition of IFNγ only during the early priming phase resulted in similar frequencies (Fig. 7f-g) and numbers (Fig. 7h) of ASCs compared to cultures that contained IFNγ throughout the entire culture period.

Excluding R848 from the cultures during the first three days (Fig. 7i) had a minimal impact on pre-ASC formation (Fig. 7j) or the proliferative history (Fig. 7k-l) and differentiation potential (Fig. 7n-o) of the cells. However, cell recovery on day 6 was significantly decreased when R848 was excluded during the priming phase (Fig. 7m), which impacted the number of ASCs recovered (Fig. 7p), suggesting that early R848 signals likely enhances cell survival during the differentiation phase. When R848 was selectively excluded in the differentiation phase, proliferation was severely stunted (Fig. 7k-l), resulting in greatly reduced cell recovery (Fig. 7m) and ASC recovery (Fig. 7p) on day 6. These data therefore indicated that R848 plays both early and late roles in the development of ASCs with early TLR7/8 signals appearing to condition the B cells to survive during the later TLR7/8-dependent proliferative phase.

When IL-21 was only included for the first three days of the culture (Fig. 7q), pre-ASCs formed normally (Fig. 7r) but proliferation (Fig. 7s-t) and cell recovery (Fig. 7u) on day 6 was extremely low. This resulted in an inability to detect (Fig. 7v-w) and recover ASCs on day 6 (Fig. 7x). By contrast, the proliferation (Fig. 7s-t) and recovery (Fig. 7u) of cells stimulated with IL-21 only during the late expansion/differentiation phase were not significantly different from cells that were stimulated for all six days in the presence of IL-21. Moreover, the frequency (Fig. 7v-w) and number (Fig. 7x) of ASCs recovered on day 6 were very similar, indicating that late IL-21 signals are sufficient to drive ASC formation. Collectively, the data show that while inclusion of IFNγ, TLR7/8 ligand and IL-21 throughout the entire culture period promotes optimal ASC recovery, the window in which each stimulus was required was distinct with IFNγ playing an important role in the priming phase, IL-21 signals contributing during the later expansion and differentiation phase of the response and R848 being important throughout the culture period (Fig. 7y). Importantly, the data also indicated that the combination of R848 and IL-21, even when included for all 6 days of the culture was not sufficient to induce ASC formation and recovery. Thus, IFNγ signals appear to synergize with R848 and IL-21 signals to promote ASC formation.

### IFNγ synergizes with IL-2 to promote ASC recovery

Our data indicated that IFNγ signals play a non-redundant and critical role in the formation of the Tbet^hi^IRF4^int^ B_DN_ pre-ASC population *in vitro*, and is necessary for both the development and recovery of ASCs in the cultures, even when IL-21 and R848 are present in the cultures. These data led us to hypothesize that IFNγ signaling might program B cells to optimally respond to stimuli, like IL-21, IL-2 and TLR ligands, that promote B cell proliferation and differentiation. To test this hypothesis, we first addressed whether IFNγ cooperates with IL-2 to promote ASC development. We activated HD B_N_ cells with anti-Ig + R848 and then washed and incubated the cells for an additional 3 days with IL-21 + R848 (Be.0 conditions; Fig. 8a). Alternatively, we added IL-2 (Be.IL2), IFNγ (Be.IFNγ conditions) or both IL-2 and IFNγ (Be.γ2 conditions) to the cultures during the first three days (Fig. 8a) and then stimulated the cells for an additional 3 days with R848 + IL-21. As expected, from our earlier experiments, we recovered very few cells and no ASCs from the Be.0 cells on day 6 (data not shown). When we examined the B cells from the Be.IFNγ, Be.IL2 and Be.γ2 cultures we observed that the B cells that were exposed to IFNγ during the first 3 days proliferated more, regardless of whether we looked on day 3 (Fig. 8b-c) or day 6 (Fig. 8d-e). This effect was relatively modest at day 3 as 20% of the Be.IL2 cells vs 42-53% of the Be.IFNγ and Be.γ2 cells had divided once over 3 days (Fig. 8b-c). Consistent with this, cell recovery on day 3 was only reduced by 20% in the Be.IL2 cultures (Fig. 8f). However, by day 6 we observed a 4-fold reduction in cell recovery between the Be.IL2 and Be.IFNγ cultures (Fig. 8g) and a 7-fold reduction in cell recovery between Be.IL2 and Be.γ2 cultures (Fig. 8g). Moreover, while the fraction of ASCs present in all 3 cultures were not significantly different (Fig. 8h-i), the number of ASCs recovered in the Be.IL2 cultures was significantly lower (Fig. 8j) when compared to either the Be.IFNγ or Be.γ2 cultures. Interestingly, the recovery of ASCs on day 6 was significantly better in the Be.γ2 cultures compared to the Be.IFNγ or Be.IL2 cultures (Fig. 8j). Thus, while early IL-2 and IFNγ signals promote cell cycle entry by BCR and TLR7/8 activated B cells, only early IFNγ signals effectively sustained the proliferative and differentiation potential of the B cells. Finally, since the combined total of ASCs recovered from the Be.IFNγ and Be.IL2 cultures was less than that recovered from the Be.γ2 cultures, the data suggested that IFNγ likely cooperates with IL-2 to promote ASC recovery.

**Figure 8.**
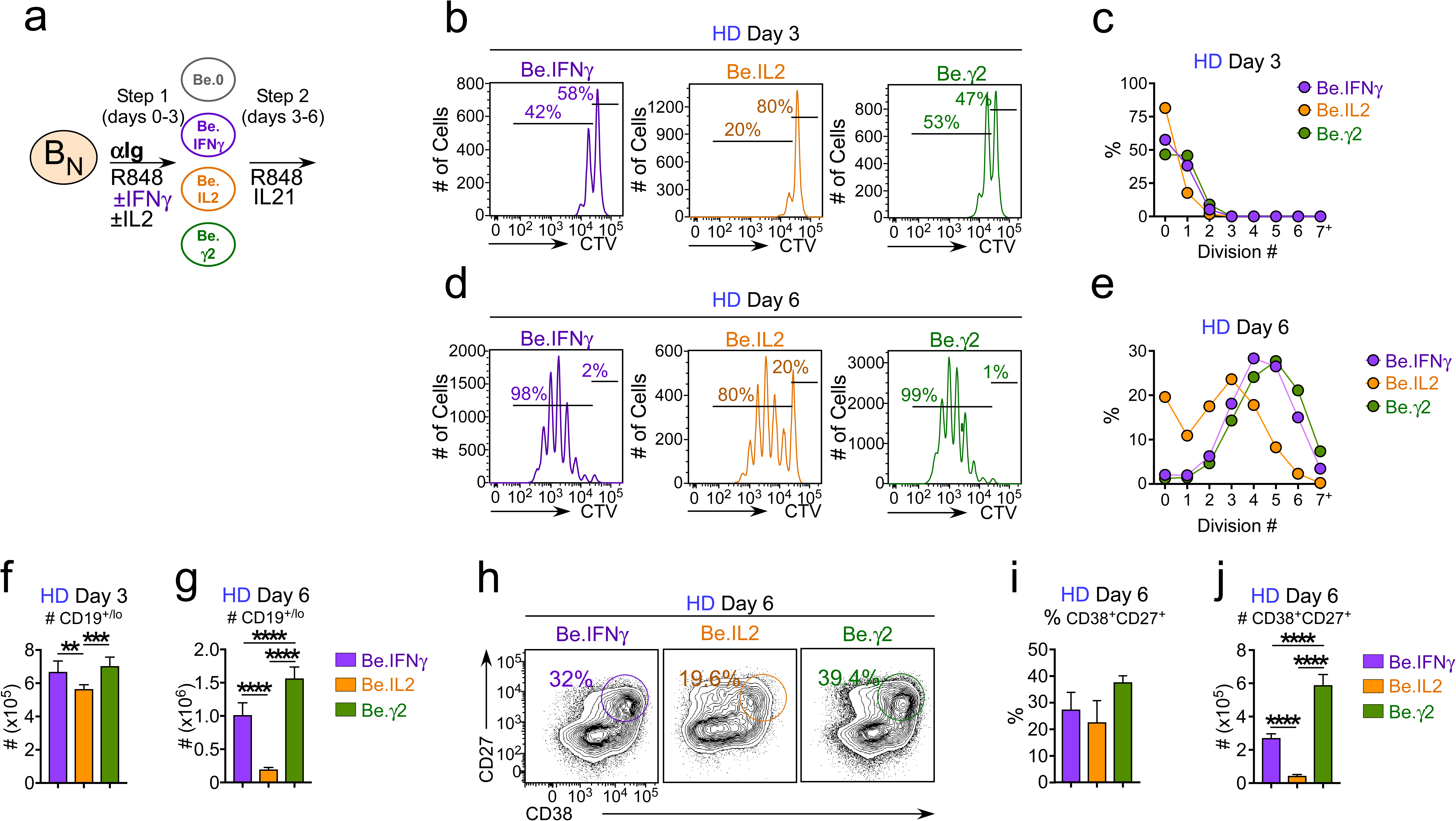
IL-2 signals cooperate with IFNγ to enhance ASC recovery from stimulated B_N_ cells. Cartoon (**a**) depicting CTV-labeled HD B_N_ cell cultures activated for 3 days (Step 1) with anti-Ig and R848 alone (Be.0) or in combination with: IFNγ (Be.IFNγ), IL-2 (Be.IL2) or both IFNγ + IL-2 (Be.γ2) and then washed and recultured for an additional 3 days (Step 2) with R848 and IL-21. Representative CTV dilution plots and enumeration of cell division within the cultures on days 3 (**b-c**) and 6 (**d-e**). Total cell recovery on days 3 (**f**) and 6 (**g**). ASC frequencies (**h-i**) and ASC recovery (**j**) in day 6 cultures. Flow cytometry plots depicting CTV dilution and CD27^hi^CD38^hi^ ASCs in each culture are representative of ≥3 experiments. The percentage of cells in each division, the frequency of ASCs and cell recovery (total and ASCs) are shown as the mean ± SD of cultures containing purified B_N_ cells from 2 independent healthy donors. All statistical analyses were performed using one-way ANOVA with Tukey’s multiple comparison test. *P* values *<0.05, **<0.01, ***<0.001, ****<0.0001.

### IFNγ synergizes with subthreshold TLR7/8 signals to promote ASC recovery

Our earlier upstream regulator analysis of T-bet^hi^ B_DN_ cells from Th1 co-cultures indicated that the TLR signaling pathway was activated in these cells (Fig. 5a), even though we did not add TLR ligands to the cultures. This suggested that the T-bet^hi^ B_DN_ cells, which were activated in the presence of IFNγ-producing Th1 cells, might be more responsive to the low levels of endogenous TLR ligands that are released by dying cells in culture (52). To test this possibility, we activated CTV-labeled HD B_N_ cells with anti-Ig, IL-2 and increasing doses of R848 (0-10 μg/ml) in the presence and absence of IFNγ (10 ng/ml) for 3 days. We then washed the cells and re-cultured the cells for an additional 3 days with IL-21 and the same concentration of R848 that the cells were exposed to during the priming phase. On day 6 we measured cell division and ASC formation. Consistent with our earlier experiments (Fig. 7), when R848 was not included in the cultures during the first 3 days the cells remained largely undivided even out to day 6 (Fig. 9a) and few ASCs were detected (Fig. 9b), regardless of whether the cells were exposed to IFNγ during first three days. By contrast, when high dose R848 was included in the first three days, the cells proliferated efficiently with or without IFNγ (Fig. 9a), although the presence of IFNγ did increase the fraction of cells that had undergone 6+ cell divisions (Fig. 9a). Interestingly, when we activated B_N_ cells with a 10-fold lower dose of TLR7/8 ligand (0.1 μg/ml), we observed minimal proliferation in the cultures that did not include IFNγ (≤3% divided ≥3 times) versus robust proliferation (63% divided ≥3 times) in the cultures that contained IFNγ (Fig. 9a). More striking, we observed that the frequency of ASCs in the cultures that were activated with low dose TLR ligand in the presence of IFNγ was approximately 10-fold higher than that observed for the cultures that lacked IFNγ (Fig. 9b). These data clearly demonstrate that exposure of B_N_ cells IFNγ during the initial priming phase allowed these cells to respond to a sub-optimal dose of R848 and to differentiate into ASCs.

**Figure 9.**
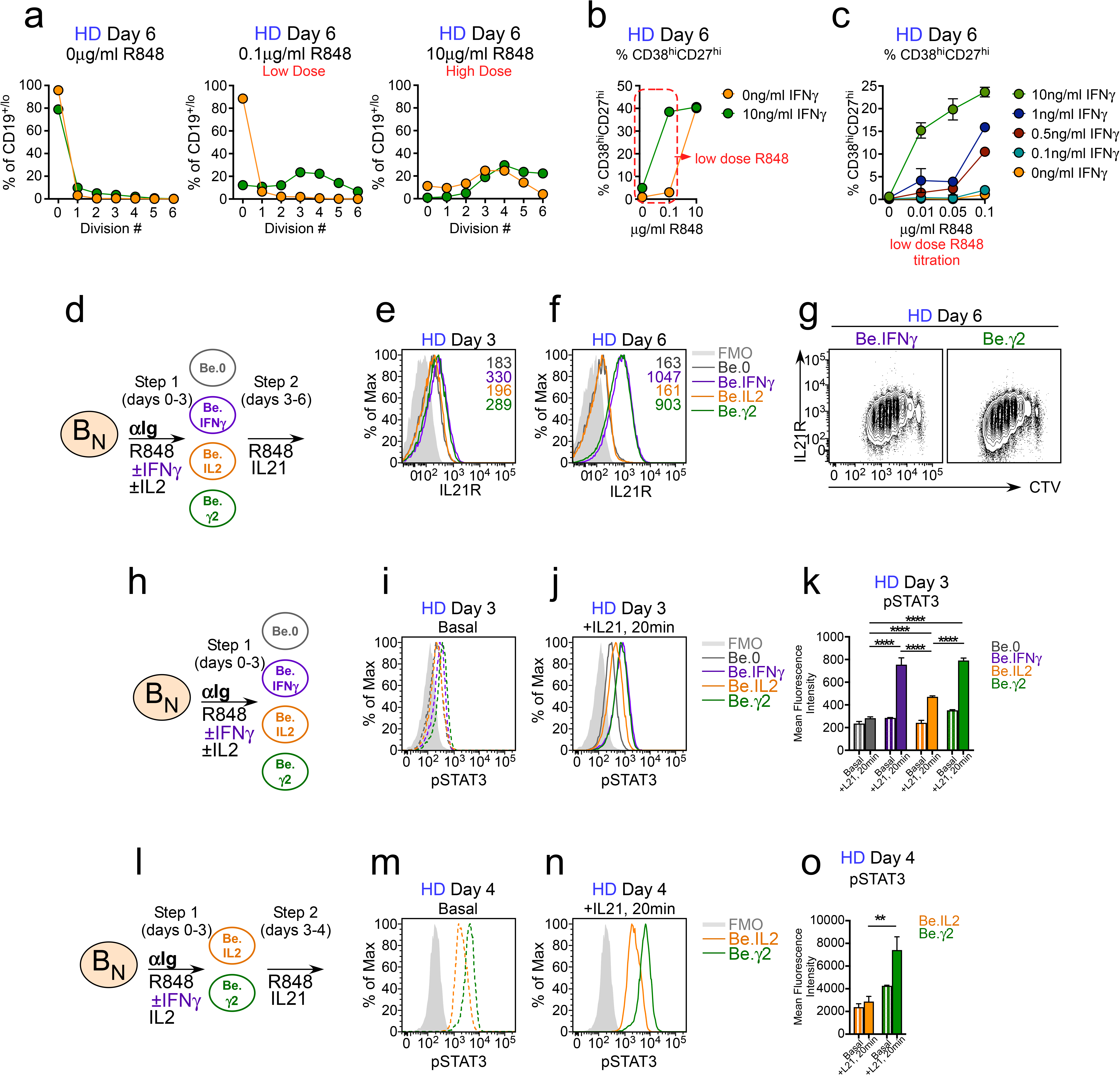
IFNγ signaling enhances B cell responses to TLR7/8 ligands and IL-21. (**a-c**) IFNγ synergizes with subthreshold doses of TLR7/8 ligand to induce proliferation and differentiation of B_N_ cells. CTV-labeled HD B_N_ cells were activated for 3 days (Step 1) with anti-Ig, IL-2, and R848 in the presence or absence of IFNγ (10ng/ml). Cells were washed and re-cultured for 3 additional days (Step 2) with IL-21 and the same concentration of R848 that was used in Step 1. B cell division on day 6 (**a**) in cultures that were activated with (green circles) or without (orange circles) IFNγ in the presence of no R848 (0μg/ml, **left panel**), low dose R848 (0.1 μg/ml, **center panel**) and high dose R848 (10 μg/ml, **right panel**). The frequency of CD38^hi^CD27^hi^ ASCs (**b**) present in each culture condition on day 6 is shown. (**c**) CTV-labeled naïve B cells were activated for 3 days with anti-Ig, IL-2 and normally non-stimulatory doses of R848 (0-0.1μg/ml) in combination with different concentrations of IFNγ (0-10ng/ml). Cells were washed and re-cultured for 3 additional days with IL-21 and the same concentration of R848 used in Step 1. The frequency of ASCs in the indicated cultures on day 6 is reported. (**d-g**) Early IFNγ signals control IL-21R expression levels. Cartoon (**d**) depicting culture conditions to generate Be.0, Be.IFNγ, Be.IL2 and Be.γ2 cells from CTV-labeled HD B_N_ cells. IL-21R expression was measured on day 3 (**e**) and day 6 (**f-g**) and reported as MFI. (**h-o**) Early IFNγ signals control IL-21R signaling. Day 3 (**i-k**) or Day 4 (**m-o**) Be.0, Be.IFNγ, Be.IL2 and Be.γ2 cells were generated from HD B_N_ cells as shown in the cartoons (**h,l**). Representative pSTAT3 expression levels in cells under basal conditions (no restimulation, **i,m**) or following 20 min IL-21 restimulation (**j,m**). Data are reported as Mean Fluorescence Intensity of pSTAT3 on day 3 (**k**) and day 4 (**o**). Flow cytometry plots depicting IL21R expression and pSTAT3 levels in B cells from the indicated cultures are representative of ≥2 experiments. Data in **a-c** are representative of 2 independent experiments with (**c**) showing the mean ± SD of experimental replicates. Data in **o** and **k** are from 2 (**o**) or 3 (**k**) independent donors and are shown as mean ± SD. Statistical analyses were performed using one-way ANOVA with Tukey’s multiple comparison test (**k,o**).

To test whether the combination of TLR ligands and IFNγ synergistically promoted ASC formation, we cultured HD B_N_ cells with anti-Ig + IL-2 while cross-titrating in IFNγ and R848. On day 3, we washed and recultured the cells for an additional 3 days with IL-21 plus the same concentration of R848 that was used in the priming phase. We then measured ASC formation in the cultures on day 6. As expected, when R848 was not included in the cultures, the frequency of ASC in the cultures was very low, regardless of the dose of IFNγ included during the priming phase (Fig. 9c). However, when IFNγ was present at 10 ng/ml in the priming phase, the B cells responded to concentrations of R848 that were 100-1000 times lower (0.01 μg/ml) than what is normally used to stimulate B cells (Fig. 9c). Moreover, when we examined the response of the B cells that were activated in the presence of a sub-optimal concentration of R848 (0.1 μg/ml), we observed a clear dose response to the IFNγ, with increasing concentrations of IFNγ giving rise to a higher frequency of ASCs in the cultures (Fig. 9c). These data therefore show that IFNγ signaling confers human B cells with the capacity to respond to extremely low subthreshold concentrations of TLR7/8 ligands, which when combined with IL-21 signals results in their robust proliferation and differentiation into ASCs.

### IFNγ programs IL-21R expression and responsiveness

Our data showed that early IFNγ signals synergize with TLR7/8 and IL-2 signals to poise the BCR-activated B cells to respond to proliferation and differentiation cues provided by subsequent exposure to IL-21 and R848. Given the important role for IL-21 in the commitment of human B cells to the ASC lineage (45, 46), we hypothesized that early IFNγ signals might regulate IL-21R expression and/or IL-21R signaling in the stimulated B cells. To test this hypothesis, we first measured IL-21R expression levels by day 3 and day 6 Be.0, Be.IL2, Be.IFNγ and Be.γ2 cells (Fig. 9d). Although day 3 B cells from Be.IFNγ and Be.γ2 cultures expressed slightly higher levels of IL-21R compared to B cells from Be.0 and Be.IL2 cultures (Fig. 9e), IL-21R expression was low in all groups at this timepoint. However, by day 6, IL-21R expression levels were significantly increased in the Be.IFNγ and Be.γ2 cells relative to the Be.0 and Be.IL2 cells (Fig. 9f). Upregulation of the IL-21R by B cells from the Be.IFNγ and Be.γ2 cultures was not linked to cell division as even the undivided cells in these day 6 cultures expressed high levels of IL-21R (Fig. 9g). Moreover, IL-21R expression levels were directly comparable between day 6 Be.IFNγ and Be.γ2 cells (Fig. 9f), demonstrating that early IFNγ signals were necessary and sufficient to program upregulation of IL-21R.

To address whether early IFNγ stimulation also increased responsiveness to IL-21, we generated day 3 Be.0, Be.IFNγ, Be.IL2 and Be.γ2 cells (Fig. 9h) and measured phospho-STAT3 levels in these cells under basal conditions and following 20 min stimulation with IL-21. Day 3 basal levels of pSTAT3 were similar between the Be.0, Be.IL2 and Be.IFNγ cells and modestly higher in the Be.γ2 cells (Fig. 9i). However, following a 20 min IL-21 stimulation pSTAT3 levels were increased by 2-fold in the B cells that were exposed to IFNγ during the priming phase (Fig. 9j-k). More importantly, when Be.IL2 and Be.γ2 cells were washed on day 3 to remove IFNγ and IL-2 and then recultured for 24 hrs in the presence of IL-21 and R848 (Fig. 9l), basal pSTAT3 levels were increased in the Be.γ2 cells relative to the Be.IL2 cells (Fig. 9m). Similarly, pSTAT3 levels in the day 4 Be.γ2 cells were higher compared to Be.IL2 cells following a 20 min restimulation with IL-21 (Fig. 9n-o). Thus, early IFNγ stimulation, particularly when combined with IL-2, promoted increased IL21R expression and enhanced IL-21R signaling.

### Early IFNγ signals alter the epigenetic landscape of activated B_N_ cells and poise T-bet^hi^ B_DN_ cells to differentiate

To address how IFNγ signaling cooperates with IL-2 and TLR7/8 ligands to poise B cells to differentiate in response to IL-21, we considered the possibility that IFNγ signaling might alter the epigenetic profile of the T-bet^hi^ B_DN_ B cells since T-bet is known to alter chromatin accessibility by recruiting chromatin-modifying enzymes to regulatory promoter and enhancer regions (14). To address this hypothesis, we performed ATAC-seq analysis on day 3 Be.0, Be.IL2, Be.IFNγ and Be.γ2 cells and identified differentially accessible regions (DARs) that were assigned to different gene loci across the genome (Supplemental File 3). As expected, distinct sets of DARs were found in all 4 groups (Fig. 10a), however the Be.γ2 cells had the greatest number of DARs (Fig. 10a). Moreover, it appeared that the chromatin accessibility pattern in the Be.γ2 cells reflected cooperation or synergy between the IFNγ and IL-2 signals (Fig. 10a). To determine whether specific transcription factor (TF) binding motifs were differentially enriched in the DAR of the Be.IL2, Be.IFNγ and Be.γ2 genomes, enriched TF-binding motifs were identified in each cell population relative to the Be.0 cells (Supplementary File 4). Chromatin accessibility surrounding TFs binding motifs, like AP1, RUNX, PU.1 and NFAT, was greatly enriched in all 3 groups relative to the Be.0 cells (Fig. 10b). Accessibility around other TF binding motifs like, IRF1 and T-bet, was enriched specifically in the B cells that were exposed to IFNγ, while accessibility at STAT5 binding motifs was enriched in IL-2 exposed B cells (Fig. 10b, Supplementary File 4). Interestingly, despite the fact that all 4 groups, including Be.0 cells, were exposed to high dose TLR7/8 ligand and anti-Ig, we observed enrichment in NF-κB p65 and NF-κB REL binding motifs within the Be.IL2, Be.γ and Be.γ2 DAR relative to Be.0 cells (Fig. 10b). However, when we examined accessibility within the 100 bp surrounding the NF-κB binding motifs, it was clear that NF-κB binding sites were most accessible in the Be.γ2 cells compared to all other groups (Fig. 10c-d, Supplementary Files 4-5). This was also true when we examined accessibility surrounding STAT5 binding motifs and T-bet binding motifs (Fig. 10e-f, Supplementary Files 5). Thus, the combination of early IFNγ and IL-2 signals in BCR and TLR7/8-activated B_N_ cells significantly and synergistically increased chromatin accessibility surrounding T-bet, STAT5 and NF-κB binding sites within the activated B cells.

**Figure 10.**
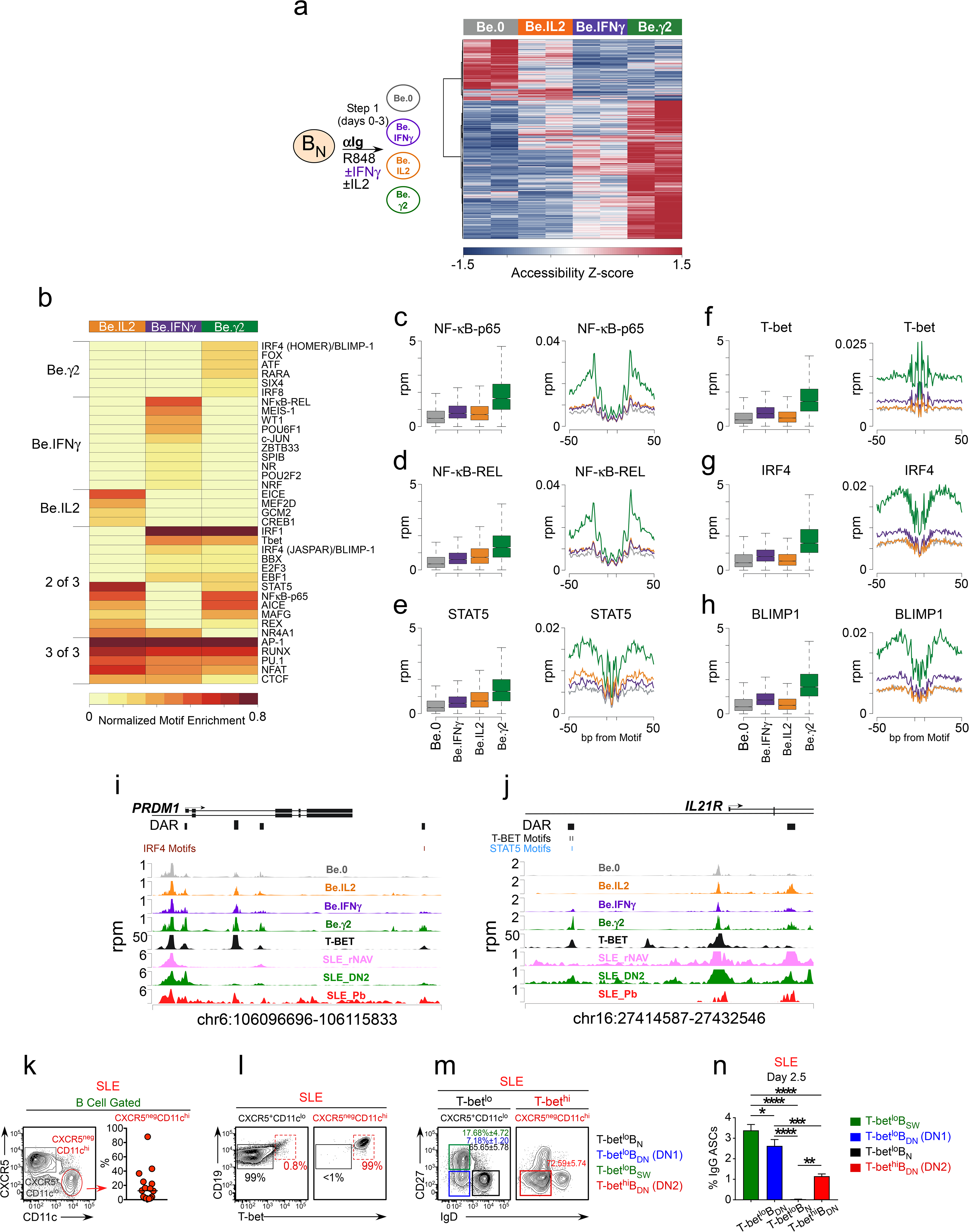
IFNγ signaling enhance chromatin accessibility and prepares BCR and TLR activated B cells to differentiate. (**a-j**) Analysis of chromatin accessibility in IFNγ-activated B cells. Cartoon (**a, left**) depicting generation of day 3 Be.0, Be.IFNγ, Be.IL2 and Be.γ2 cells from HD B_N_ cells. Heatmap (**a, right**) of ATAC-seq data (Supplementary File 3) from the four B cell subsets showing 15,917 differentially accessible regions (DAR) based on FDR<0.05. (**b**) Heatmap depicting enrichment of accessible transcription factor motifs in day 3 Be.IFNγ, Be.IL2 and Be.γ2 subsets generated from HD B_N_ cells. Data are shown as normalized motif enrichment within each subset relative to Be.0 cells. Motif grouping (indicated on left) showing transcription factors that are enriched for accessibility in 1 subset, 2 subsets or all 3 subsets. See Supplementary File 4 for complete results of Homer *de novo* Motif Results. (**c-h**) Chromatin accessibility for NF-κB p65 (**c**), NF-κB REL (**d**), STAT5 (**e**), T-bet (**f**), IRF4 (**g**) and BLIMP1 (**h**) in day 3 B cell subsets. Data represented as box and whisker plots, which report reads per million (rpm) in the 100bp surrounding the transcription factor binding motifs, or histograms, which show accessibility at the indicated motif and for the indicated surrounding sequence. *P* values provided in Supplementary File 5; Be.γ2 is significantly different from all other B cell subsets in all analyses (*P*<2.2 x10^-16^). (**i-j**) Genome plot showing chromatin accessibility for the *PRMD1* (**i**) and *IL21R* (**j**) loci in day 3 Be.0, Be.IFNγ, Be.IL2 and Be.γ2 cells generated from HD B_N_ cells. DAR are shown and consensus T-bet, IRF4 and STAT5 binding motifs within DARs are indicated. DAR are aligned with previously reported T-bet binding sites (assessed by ChIP, (56)) and with ATAC-seq data derived from B cell subsets purified from SLE patients (19). Data reported in rpm. (**k-n**) SLE patient T-bet^hi^ B_DN_ cells rapidly differentiate in ASCs. Sort purification strategy (**k-m**) to isolate SLE B cell subsets. Peripheral blood CD19^+/lo^ B lineage cells from SLE patients were subdivided into CD27^hi^CD38^hi^ ASCs and non-ASC B cells (see Fig. 1g) that were then further subdivided into CD11c^hi^CXCR5^neg^ and CD11c^lo^CXCR5^+^ cells (**k**, right panel). T-bet^hi^ B cells were highly enriched (99%) within the CD11c^hi^CXCR5^neg^ fraction (**l**) while T-bet^lo/neg^ cells were contained exclusively within the CD11c^lo^CXCR5^+^ fraction (**l**). The CD11c^hi^CXCR5^lo^ (T-bet^hi^) and CD11c^neg^CXCR5^+^ (T-bet^lo^) B cells were further subdivided (**m**) based on expression of IgD and CD27 and sort-purified as T-bet^lo^ B_N_ cells (black gate, CD11c^lo^CXCR5^+^IgD^+^CD27^neg^), T-bet^lo^ B_DN_ cells (purple gate, CD11c^lo^CXCR5^+^IgD^neg^CD27^neg^, also referred to as DN1 cells (19)), T-bet^lo^ B_SW_ cells (green gate, CD11c^lo^CXCR5^+^IgD^neg^CD27^+^) and T-bet^hi^ B_DN_ cells (red gate, CD11c^hi^CXCR5^neg^IgD^neg^CD27^neg^, also referred to as DN2 cells (19)). Purified SLE B cell subsets were stimulated with cytokines (IFNγ, IL-21, IL-2, Baff) and R848 for 2.5 days then counted and transferred to anti-IgG ELISPOT plates for 6 hrs. The frequency of IgG ASCs (**n**) derived from each B cell subset is shown. ATAC-seq analysis was performed on 3 independent experimental samples/group over 2 experiments. Box plots show 1^st^ and 3^rd^ quartile range (box) and upper and lower range (whisker) of 2 samples/group. Data reported in (**n**) are representative of 3 independent experiments using B cells sorted from 3 different SLE donors. *P* values for enrichment of transcription factor binding motifs (**c-h**) are provided in Supplementary File 5. Statistical analysis in (**n**) was performed with unpaired ANOVA on triplicate experimental replicates. *P* values *<0.05, **<0.01, ***<0.001, ****<0.0001.

Interestingly, chromatin accessibility surrounding the HOMER-defined IRF4 and BLIMP1 binding motif (55) was also enriched in the Be.γ2 cells (Fig. 10g-h, Supplementary Files 5). These data therefore suggested that these key ASC initiating TFs were already exerting epigenetic changes to the genome of the Be.γ2 cells, even before these cells were exposed to IL-21. Consistent with this finding, when we examined the *PRDM1* locus, we identified 4 DARs that were each more accessible in the Be.γ2 cells relative to the other cells (Fig. 10i). Interestingly, while none of these DAR contained a T-bet binding motif, each of these 4 DAR directly aligned with peaks previously identified in a published T-bet ChIP-seq analysis of GM12878 cells (56), suggesting that T-bet could be associated with TF complexes that bind to these regulatory regions. Moreover, 3 of the 4 PRDM1-associated DARs were also seen in T-bet^hi^ DN2 cells purified from SLE patients (Fig. 10i), indicating that these DARs are present in the pre-ASC population found in SLE patients.

Finally, given our data showing that IFNγ and IL-2 enhanced expression of IL-21R and potentiated signaling through this receptor, we examined the 2 DAR assigned to the *IL21R* locus of the day 3 cells (Fig. 10j). One of the DAR contained two putative T-bet binding motifs and was directly aligned with a T-bet ChIP-seq peak from GM12878 cells (56) (Fig. 10j). This DAR was only observed in the cells that were exposed to IFNγ and was most enriched in the Be.γ2 population. Interestingly, we identified the same DAR in the SLE patient T-bet^hi^ DN2 cells (Fig. 10j), which are reported to be highly responsive to IL-21 (19). Taken together, the data suggested that early IFNγ signals synergize with BCR, TLR and IL-2 signals to induce global changes in chromatin accessibility and promote increased TF binding at T-bet, NF-κB, STAT5, BLIMP1 and IRF4 binding sites as well as chromatin remodeling at the *PRDM1* and *IL21R* loci.

### SLE T-bet^hi^ DN2 cells differentiate rapidly into ASCs

Previous data from our group showed that the T-bet^hi^ DN2 cells from SLE patients are pre-ASCs (19) but that these cells are not memory cells and, like B_N_ cells (46), are highly dependent on IL-21 signals to differentiate. Given our data showing that the DARs found in the *PRDM1* and *IL21R* loci of the day 3 T-bet^hi^IRF4^int^-expressing Be.γ2 cells were also conserved in the T-bet^hi^ DN2 cells from SLE patients, we hypothesized that stimulation of the SLE DN2 cells with IL-21, IL-2 and TLR7/8 ligand would result in rapid differentiation into ASCs. To test this hypothesis, we enumerated IgG-producing ASCs after sort-purifying SLE patient-derived T-bet^lo^ B_N_ cells, T-bet^lo^ memory B cells (T-bet^lo^ B_DN_ (DN1 memory cells) and T-bet^lo^ B_SW_) and T-bet^hi^ DN2 cells, (Fig. 10k-m) and then stimulating the cells with R848, IFNγ, IL-21 and IL-2 for 2.5 days. As expected, the two memory B cell subsets efficiently formed ASCs in this short timeframe (Fig. 10n), while B_N_ cells failed to differentiate (Fig. 10n). However, ASCs were easily identified in the day 2.5 T-bet^hi^ DN2 cultures (Fig. 10n) and ASC recovery was at least 50-fold higher in T-bet^hi^ DN2 cell cultures compared to the B_N_ cultures and only 2-3 times less than that seen with the memory B cell populations (Fig. 10n). Thus, these data are consistent with the interpretation that the expanded population of T-bet^hi^ DN2 cells present in some SLE patients likely represent a discrete population of B_N_-derived pre-ASCs that received prior programming signals from TLR, IFNγ and antigen signals and are poised to differentiate into ASCs upon downregulation of the BCR signaling cascade and exposure to IL-21. The relevance of these findings to pathogenic and protective B cells responses are discussed.

## Discussion

Here we show that activation of human naïve B (B_N_) cells with allogeneic IFNγ-producing Th1 cells induces formation of a T-bet^hi^IRF4^int^ IgD^neg^CD27^neg^ (B_DN_) population that is transcriptionally and phenotypically very similar to the T-bet expressing CD11c^hi^CXCR5^lo^ B_DN_ (referred to as DN2 cells) subset found in SLE patients (19) and the CD11c^hi^ Age-Associated B cells (ABCs) that accumulate in aged and autoimmune mice and humans (18, 27). The *in vitro* generated T-bet^hi^IRF4^int^ B_DN_ cells are transcriptionally poised to become ASCs as they express a number of ASC-specific genes, including genes that are direct targets of the ASC commitment transcription factor IRF4. Moreover, a significant fraction (up to 40%) of sorted T-bet^hi^IRF4^int^ B_DN_ cells differentiate into ASCs following a single cell division. The presence of the T-bet^hi^IRF4^int^ B_DN_ pre-ASCs, which were found in the Th1/B_N_ co-cultures but not in Th2/B_N_ co-cultures, positively correlates with ASC formation in the cultures, suggesting that one or more signals provided by Th1 cells enhance ASC development and recovery. Our data show that one of the key Th1-derived signals that promotes ASC development is IFNγ.

Prior mouse studies provide hints that IFNγ signals can enhance ASC responses. For example, excess IFNγ produced by Tfh1-like CD4 T cells from autoimmune mice is associated with increased germinal center (GC) responses and autoAb production (57) and B cell intrinsic expression of IFNγR and the IFNγ-activated transcription factor, STAT1 (20, 21), is necessary for the development of autoAbs in some SLE models. Consistent with these results, multiple reports indicate that IFNγR signaling may be dysregulated in patients with autoAb-mediated disease, particularly those with SLE (9). For example, prior studies show that the transcriptome of PBMCs from some SLE patients is enriched in IFNγ-induced genes (10, 11) and other publications reveal that mRNA and protein levels of T-bet, a known IFNγ-induced transcription factor (14), are elevated in T (8) and B cells (17-19) from SLE patients. Moreover, patients with active disease are more likely to exhibit a skewed “Th1-like” profile (8, 16) as measured by determining the ratios of T-bet to GATA-3 transcripts or IFNγ to IL-4 transcripts. Finally, elevated serum IFNγ can be observed years before the onset of clinical disease in many patients (12, 13).

Despite all of the indirect data suggesting a role for IFNγ signaling in human B cell autoAb responses, surprisingly little has been done to address how, at a mechanistic level, IFNγ signaling might shape B cell Ab responses in either mice or humans. This is particularly true for human B cells as earlier *in vitro* experiments with IFNγ-stimulated human B cells revealed only very modest effects on activation and differentiation (58-60) and reports examining vaccine responses in STAT1 deficient patients (61, 62) indicated that these individuals were competent to produce Abs in response to at least some vaccines. These data, which argue that IFNγ and STAT1 signaling are not obligate for human B cell Ab responses, agree with our *in vitro* studies showing that B cells can differentiate in the absence of IFNγ-induced signals. In fact, the frequency of ASCs that we detect in our co-cultures containing Th2 cells and B_N_ cells plus IL-21 and IL-21 is similar (∼5%) to that reported previously for human B_N_ cells activated with CD40L, IL-2 and IL-21 (63). Our novel finding is that B cell intrinsic IFNγ signals can greatly augment B_N_ ASC responses by enhancing proliferation and differentiation induced by TLR7/8 ligands and IL-21. Indeed, we routinely recovered 4-10 fold more ASCs in the B_N_ cultures that contain IFNγ or IFNγ-producing T cells compared to cultures that lack IFNγ. Thus, IFNγ signaling has the potential to drive ASC development in settings, like autoimmunity and viral infection, where type 1 inflammatory cytokines and TLR ligands are present.

Our data show that increased recovery of ASCs in the IFNγ-containing cultures is dependent on IFNγ signals that are delivered in conjunction with IL-2 and BCR + TLR7/8 ligands during the initial activation of B_N_ cells. The co-activation of B_N_ cells with IFNγ and IL-2 + BCR + TLR7/8 ligands results in IFNγ-dependent remodeling of the chromatin and the formation of the T-bet^hi^IRF4^int^ pre-ASC subset. These early IFNγ-dependent signals are required for subsequent proliferation and differentiation following stimulation with IL-21 and TLR7/8 ligand. IFNγ is not, in and of itself, a B cell mitogen and has, in fact, been reported to induce apoptosis of human B cells (64, 65). However, we find that IFNγ synergizes with TLR7/8 ligand signals to promote multiple rounds of B cell proliferation – an important prerequisite of ASC differentiation (66). Although *in vitro* experiments using human B cells show that IFNγ can synergize with TLR7 and CD40 signals to promote upregulation of Bcl6 and the acquisition of a germinal center-like phenotype (21), there are no reports of IFNγ and TLR7/8 signals cooperating to promote human B cell proliferation or differentiation into ASCs. By contrast, it is well appreciated that IFNα-directed signals can enhance TLR7-mediated human B cell proliferation and differentiation (67, 68). Given that there is considerable overlap between genes regulated by IFNα and IFNγ (9), it is possible that IFNα and IFNγ may augment TLR7 signaling in human B cells by similar mechanisms.

Our *in vitro* data suggest at least three ways in which early IFNγ priming signals promote subsequent IL-21-dependent ASC differentiation. First, we show that IFNγ signals, particularly when combined with IL-2, promote global epigenetic alterations in chromatin accessibility of BCR + TLR7/8 ligand activated B_N_ cells. These epigenetic changes are associated with increased chromatin accessibility surrounding NF-κB, STAT5 and T-bet binding sites. While it is not particularly surprising that IFNγ signaling induces increased T-bet expression and alterations in chromatin accessibility around T-bet binding sites (see e.g. (69)), the finding that accessibility surrounding NF-κB and STAT5 binding motifs is also initiated in an IFNγ-dependent manner in B cells suggests that IFNγ likely augments TLR-and IL-2-dependent activation of NF-κB and STAT5, thereby allowing these transcription factors to reshape the B cell epigenome to favor robust B cell proliferation and differentiation. This result is consistent with data examining CD8 T cells that showed that IL-2R/STAT5 signaling is important for expression of BLIMP1 and that BLIMP1 and T-bet cooperate to induce effector cell differentiation (70). Second, we show that IFNγ promotes commitment to the ASC lineage by regulating two key ASC transcription factors (41), IRF4 and BLIMP1. Indeed, our data suggest that IFNγ promotes remodeling of the B cell epigenome to an ASC permissive state by: (i) regulating chromatin accessibility surrounding IRF4 and BLIMP1 binding sites within regulatory regions in the genome of the activated B_N_ cells; (ii) promoting chromatin accessibility within the *PRDM1* (BLIMP1) locus; and (iii) inducing IRF4 expression by the activated B_N_ cells. Finally, we demonstrate that IFNγ signals alter chromatin accessibility within the *IL21R* locus of the activated B_N_ cells and that this change in accessibility is associated with IFNγ-dependent, increased expression of the IL-21R by the activated B_N_ cells and with increased responsiveness of these cells to IL-21 as measured by phosphorylation of STAT3. While the important role for IL-21 in human B cell differentiation is well appreciated (45, 46), this is the first time, to our knowledge, that IFNγ signals have been shown to poise human B_N_ cells to respond to subsequent IL-21 signals.

We show that IFNγ signaling is necessary for upregulation of T-bet by B_N_ cells and changes in chromatin accessibility surrounding T-bet binding motifs. Given the known role for T-bet in controlling the recruitment of chromatin modifying enzymes like Jmjd3 H3K27 demethylases and Set7/9 H3K4 methyltransferases (14) to DNA, it is possible that the IFNγ-induced changes in chromatin accessibility, may be due, at least in part, to T-bet. Importantly, T-bet binding motifs are found in the IFNγ-dependent differentially accessible region (DAR) that is present in the *IL21R* locus of IFNγ-activated human B cells. Moreover, this IFNγ-dependent, T-bet motif containing DAR in the *IL21R* locus maps to a known site of T-bet binding in B cells, as determined by T-bet ChIP-seq (56). Similarly, the DARs that we identified in the *PRDM1* locus of the BCR, TLR7/8, IFNγ and IL-2 activated B_N_ cells also align with T-bet binding sites identified by ChIP-seq in B cells (56). While it is tempting to speculate that T-bet may directly promote expression of *PRDM1*, it is reported that IL-2R-activated STAT5, and not T-bet, which is responsible for inducing BLIMP1 in CD8 T cells (70). Therefore, it is possible that upregulation of *PRDM1* in B cells is also controlled by STAT5 and that T-bet’s key role is in regulating *IL21R* expression levels.

The IFNγ-induced T-bet^hi^IRF4^int^ pre-ASC population that we identified in our *in vitro* studies is very similar at phenotypic, molecular and functional levels to the T-bet^hi^ DN2 subset that is expanded in SLE patients. Since the expansion of the T-bet^hi^ DN2 cells in SLE patients correlates with systemic levels of IFNγ and IFNγ-induced cytokines, we think it likely that these cells arise in an IFNγ-dependent fashion in the patients and more importantly, that these cells have undergone IFNγ-dependent epigenetic programming. In support of this possibility, we demonstrate that the IFNγ-directed changes in chromatin accessibility within the *IL21R* and *PRDM1* loci seen in the *in vitro* generated T-bet^hi^IRF4^int^ pre-ASCs are also found in T-bet^hi^ DN2 cells isolated from SLE patients. Moreover, we show that B_N_ cells from SLE patients can, when activated in the presence of BCR and TLR ligands plus IFNγ and IL-2, develop into pre-ASCs that are very similar to the DN2 subset. Together, these results suggest that the SLE T-bet^hi^ DN2 cells are likely to represent antigen-activated, IFNγ-programmed primary effectors rather than atypical memory B cells (29). This hypothesis is consistent with published data (18, 19) showing that T-bet^hi^ DN2 cells, like B_N_ cells (71), require IL-21-driven STAT3 signals to differentiate and with our data showing that SLE patient T-bet^hi^ DN2 cells, unlike B_N_ cells, rapidly differentiate into ASCs in the absence of BCR signaling. Thus, our data suggest a model in which B_N_ cells from SLE patients are activated in the autoimmune microenvironment by autoantigen, endogenous TLR ligands and IFNγ and then go on to receive IL-21 differentiation signals, presumably delivered by T cells. While these interactions could take place in primary germinal center reactions, our results are also consistent with data from mouse SLE models showing that autoAb-producing ASCs can develop in a TLR-dependent and T cell-driven manner outside of the germinal center within extrafollicular sites (72). Intriguingly, recent data from autoimmune patients show that IL-21 and IFNγ co-producing Th cells (TpH cells) are localized in the perifollicular region of secondary lymphoid tissues in autoimmune patients (73).

One of our key findings is that IFNγ-augmented ASC formation and recovery is highly reliant on TLR7/8 activation by its RNA and RNA/protein ligands, which are derived from viral pathogens and dead and dying cells (3). Signaling through TLR7 is known to be important in SLE as prior studies reveal that SNPs in the human *TLR7* locus (74) and overexpression of TLR7 in mice (75) are associated with increased SLE susceptibility while deletion of TLR7 protects mice from the development of SLE (76). Our data show that deletion of the IFNγ-inducible transcription factor T-bet in B lineage cells prevents autoAb responses in a mouse model (75) of TLR7-dependent SLE. Moreover, our data shows that B cell intrinsic IFNγ signaling induces a TLR7/8 hyperresponsive state in human B cells. This finding did not appear to be due to IFNγ-dependent changes in the expression of TLR7 by the B cells (data not shown). Rather, we found that IFNγ-exposed B_N_ cells can respond and differentiate into ASCs when exposed to 100-fold lower concentration of TLR7/8 ligands than normally used to activate B cells. Given that we observed that even low levels of IFNγ are sufficient to synergize with suboptimal concentrations of TLR7/8 ligands, we predict that B cells from autoimmune patients with detectable systemic levels IFNγ will be highly sensitive to the presence of endogenous and exogenously derived TLR7 ligands. In support of this hypothesis, phosphorylation of MapK and Erk is increased in TLR7/8-stimulated SLE T-bet^hi^ DN2 cells (19) and we found that even relatively low concentrations of IFNγ in our co-cultures containing Th2 and B_N_ cells was sufficient to induce increased ASC recovery in the cultures (not shown). Thus, we predict that TLR7-driven ASC responses are likely to be exacerbated in individuals with IFNγ-associated inflammatory disease.

Our data showing that IFNγ enhances TLR7/8 signaling in B cells suggest that SLE patients who have higher systemic levels of IFNγ and expansion of the T-bet^hi^ DN2 population are likely to have more severe disease. Consistent with this, we (Fig. 1 and (19)) and others (18) show that the size of this population correlates with autoAb titers and with disease activity in SLE patients. Interestingly, prior reports demonstrate that the T-bet^hi^ DN2 cells and T-bet^hi^ CD11c^hi^ ABCs are most expanded in African American SLE patients and in patients with higher disease activity (18, 19). Similarly, we found that the patients in our cohort with the most elevated levels of IFNγ and the largest expansion of the T-bet^hi^ DN2 cells were all African American while Caucasian patients were uniformly low for both circulating T-bet^hi^ DN2 cells and systemic IFNγ (data not shown). However, it is important to note that T-bet^hi^ DN2 cells are unlikely to represent a purely “pathogenic” population as we also identified an inducible population of vaccine-specific T-bet^hi^ DN2 cells in healthy individuals who were immunized with inactivated influenza virus (data not shown). Similarly, others have reported a T-bet expressing effector <u>switched memory</u> subset (CD27^+^CD21^lo^) with pre-ASC attributes, which is induced following vaccination or infection (33, 40). Thus, we speculate that the T-bet^hi^ primary effector and memory B cell subsets, which are found in HD and autoimmune patients in the settings of acute and chronic inflammation driven by vaccination, infection, autoimmunity and aging, are formed in an IFNγ-dependent manner and likely represent a pool of pre-ASCs that are poised to differentiate.

In summary, we demonstrate that IFNγ is critical for the formation of a T-bet^hi^IRF4^int^ pre-ASC population that is remarkably similar to the T-bet^hi^ DN2 cells that accumulate in SLE patients who present with high autoAb titers, elevated disease activity and increased systemic levels of IFNγ. We show that IFNγ signals, particularly when combined with IL-2 and TLR7/8 + BCR ligands, initiate epigenetic reprogramming of human B cells – changes which poise the activated B_N_ cells to respond to IL-21 and fully commit to the ASC lineage. Based on these results, we predict that blocking IFNγ signaling in SLE patients should curtail development of T-bet^hi^ DN2 pre-ASCs, which may result in decreased autoAb production and reduced disease activity. Interestingly, and in opposition to our hypothesis, results from a phase I trial examining IFNγ blockade in SLE patients did not reveal a therapeutic benefit (77). However, no African Americans SLE patients with nephritis were included in the study (77) and, given the data showing that T-bet^hi^ DN2 cells are most expanded in African American patients with severe disease (18, 19) and our data suggesting that these cells develop in an IFNγ-dependent manner, we propose that future studies evaluating the efficacy of IFNγ blockade in SLE patients should be focused specifically on those patients who present with elevated IFNγ levels and significant expansion of the T-bet expressing DN2 pre-ASC population.

## Materials and Methods

### Key Resources Table

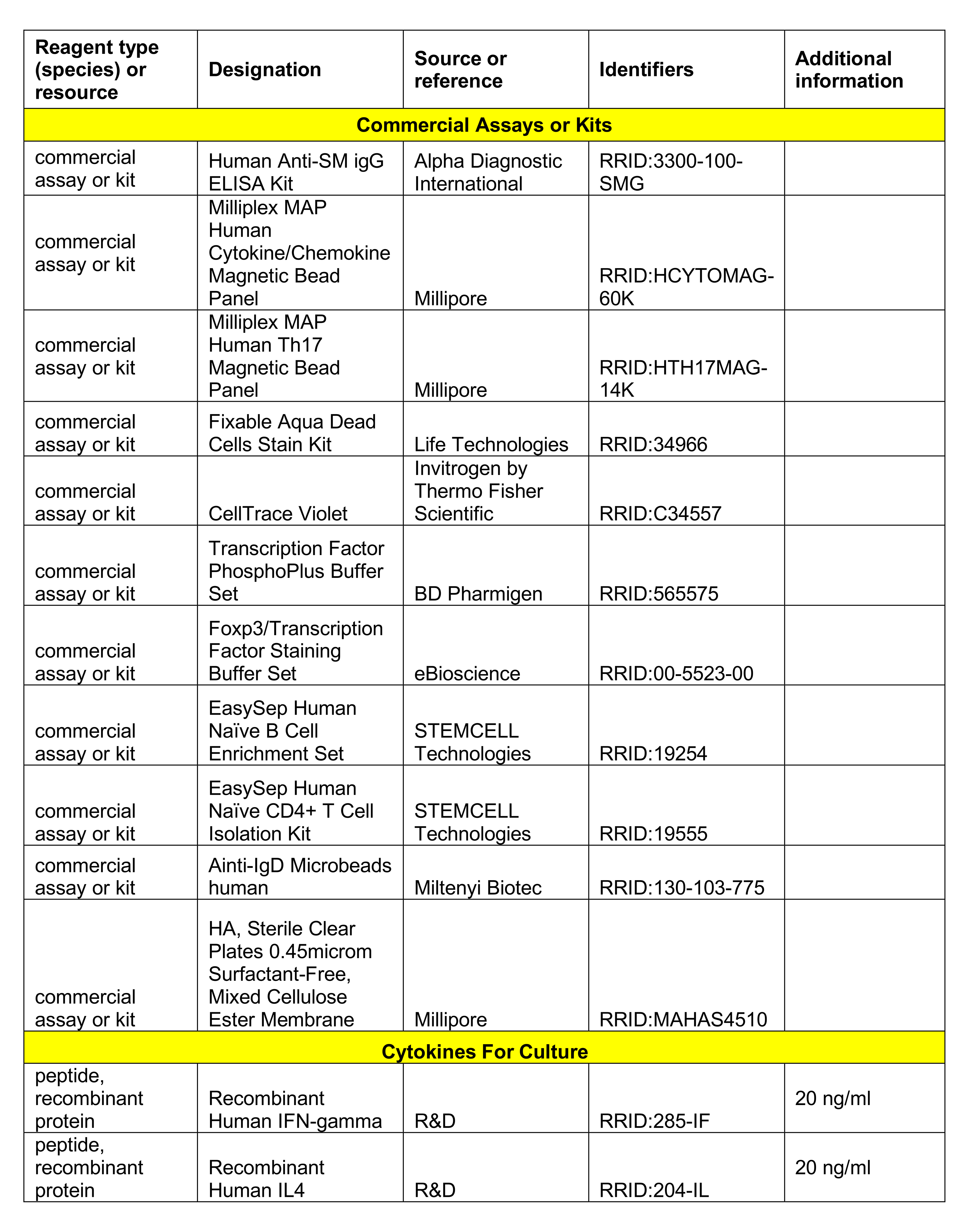
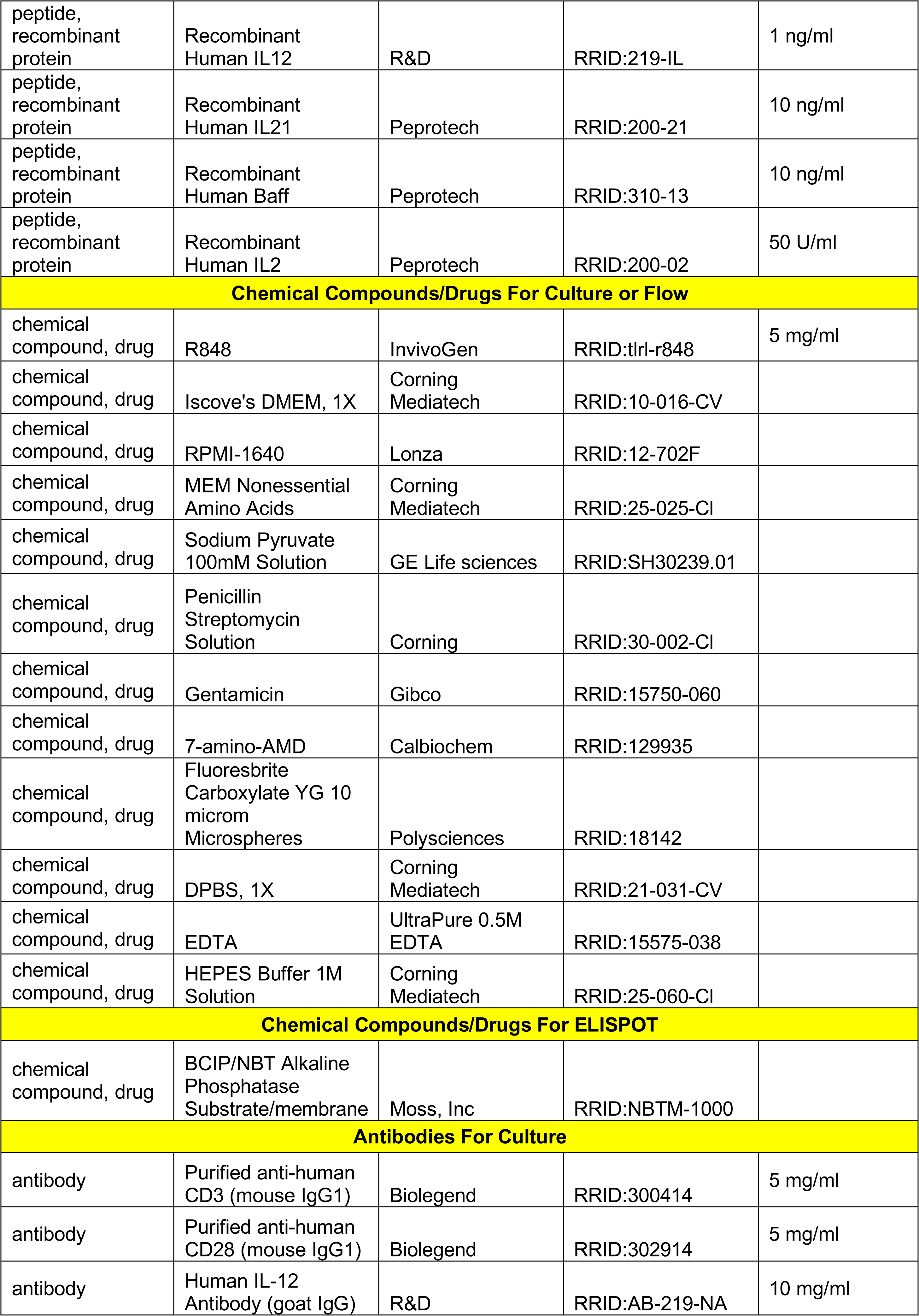
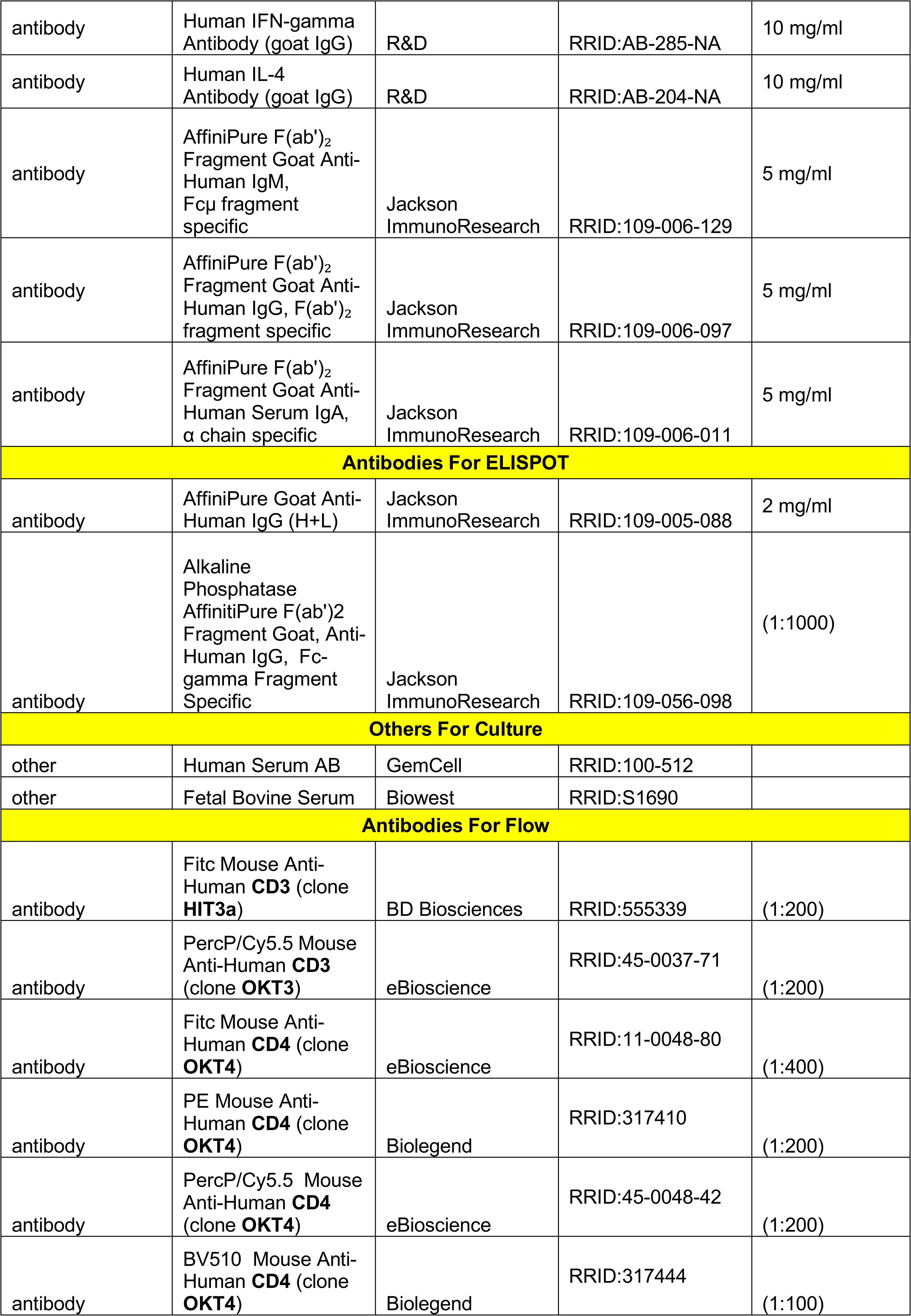
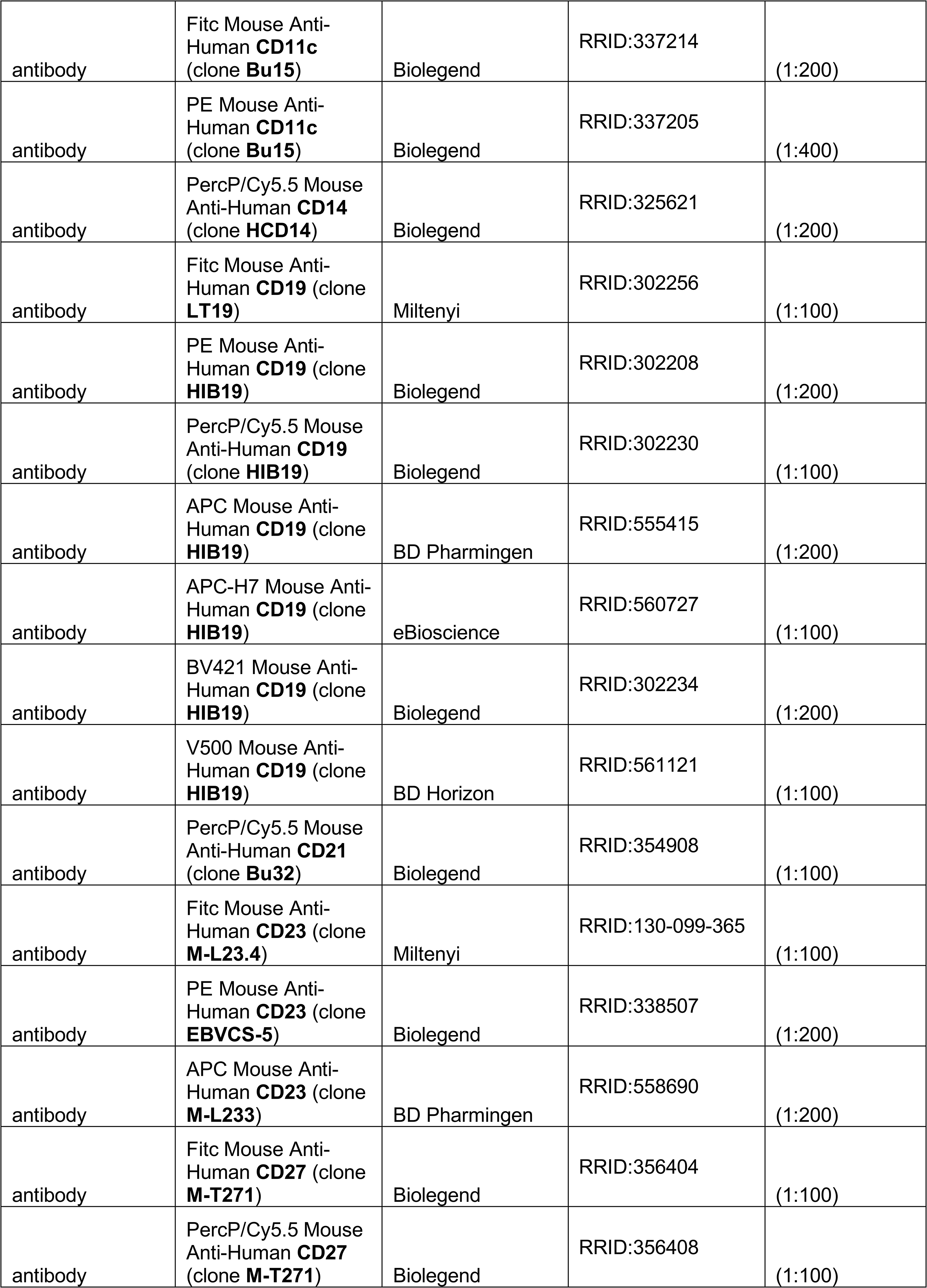
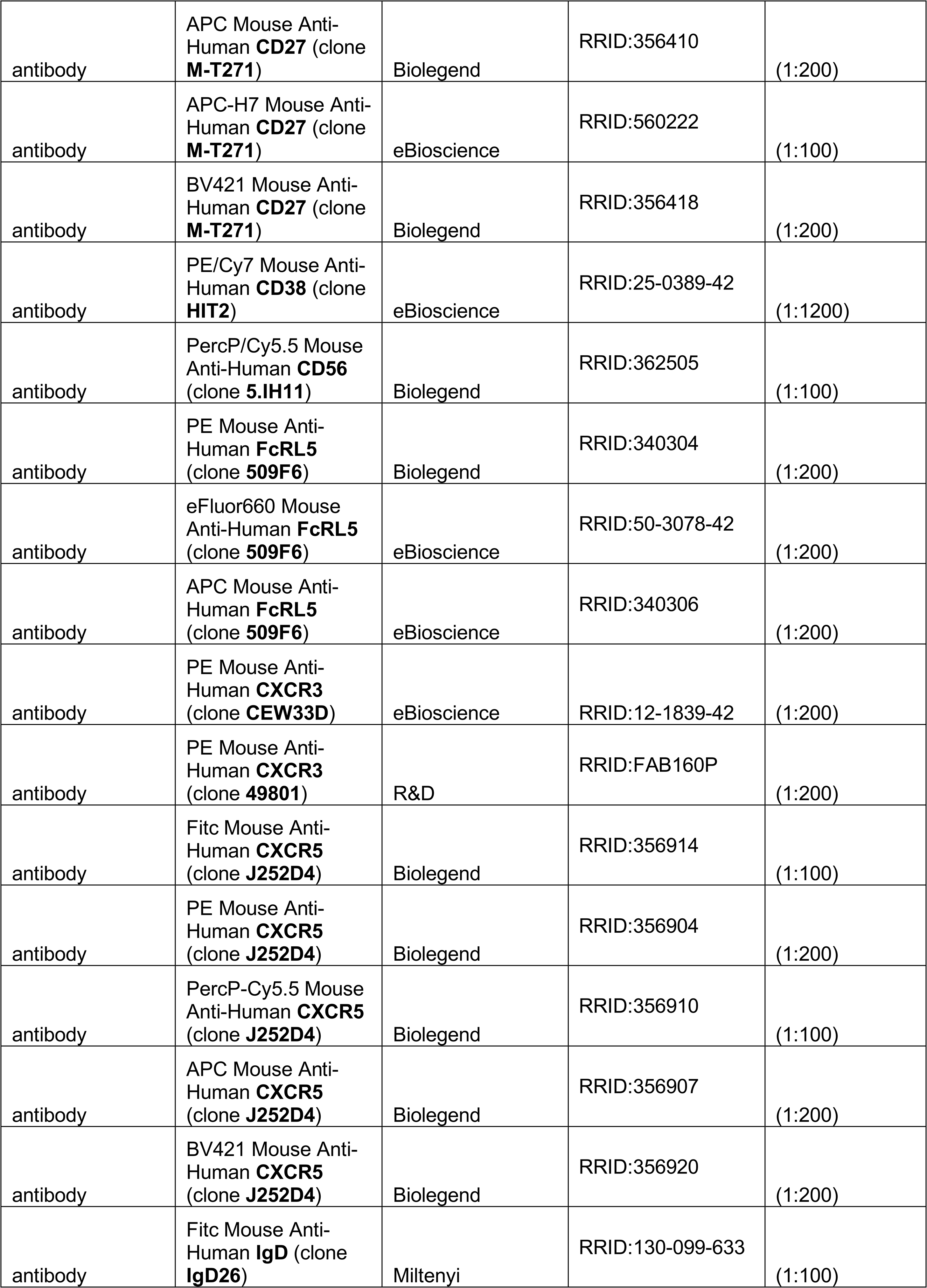
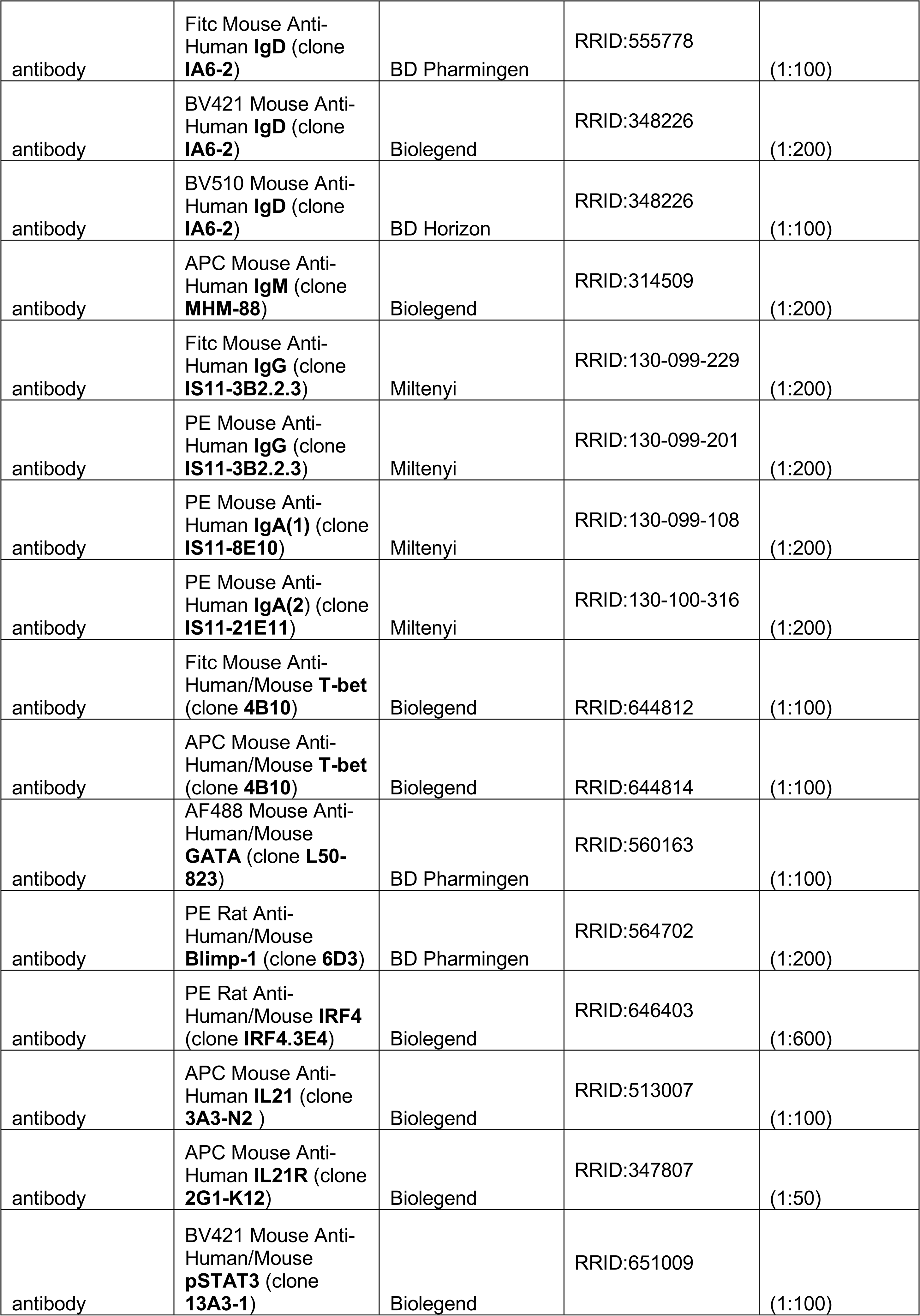

#### Human Subjects and samples

The UAB and Emory Human Subjects Institutional Review Board approved all study protocols for HD (UAB) and SLE patients (UAB and Emory). All subjects gave written informed consent for participation and provided peripheral blood for analysis. SLE patients were recruited in collaboration with the outpatient facilities of the Division of Rheumatology and Clinical Immunology at UAB or the Division of Rheumatology at Emory. UAB and Emory SLE patients met a minimum of three ACR criteria for the classification of SLE. HDs were self-identified and recruited through the UAB Center for Clinical and Translational Science and the Alabama Vaccine Research Center (AVCR). The UAB Comprehensive Cancer Center Tissue Procurement Core Facility provided remnant tonsil tissue samples from patients undergoing routine tonsillectomies.

#### Lymphocyte and plasma isolation

Peripheral blood (PB) from human subjects was collected in K2-EDTA tubes (BD Bioscience). Human tonsil tissue was dissected, digested for 30 min at 37°C with DNAse (150 U/ml, Sigma) and collagenase (1.25 mg/ml, Sigma), and then passed through a 70μm cell strainer (Falcon). Human PBMCs and plasma from blood samples and low-density tonsil mononuclear cells were separated by density gradient centrifugation over Lymphocyte Separation Medium (CellGro). Red blood cells were lysed with Ammonium Chloride Solution (StemCell). Plasma was fractionated in aliquots and stored at −80°C. Human PBMCs and tonsil mononuclear cells were either used immediately or were cryopreserved at −150°C.

#### Human lymphocyte purification

Naïve CD4^+^ T cells and CD19^+^ B cells were isolated from human PBMCs or tonsils using EasySep™ enrichment kits (StemCell). B_N_ cells were then positively selected using anti-IgD microbeads (Miltenyi). B cell subsets were sort-purified from PBMCs and tonsils as described in text.

#### Generation of Th1 and Th2 cells

Polarized CD4^+^ effector T cells were generated by activating purified HD naïve CD4 T cells with plate-bound anti-CD3 (UCHT1) and anti-CD28 (CD28.2) (both 5μg/ml, Biolegend) in the presence of IL-2 (50U/ml), IL-12 (1ng/ml) and anti-IL4 (10μg/ml) (Th1 conditions) or IL-2 (50U/ml), IL-4 (20ng/ml), anti-IL12 (10μg/ml) and anti-IFNγ (10μg/ml) (Th2 conditions). Cells were transferred into fresh media on day 3 and IL-2 was added, as needed. Cells were re-activated every 7 days using the same cultures conditions for 3 rounds of polarization. All cytokines and Abs except IL-2 (Peprotech) were purchased from R&D and T cell polarizing media contained Iscove’s DMEM supplemented with penicillin (200μg/ml), streptomycin (200μg/ml), gentamicin (40μg/ml), 10% FBS and 5% human serum blood type AB.

#### T/B co-cultures

Purified B cell subsets from HD or SLE patients were co-cultured in B cell media in the presence of IL-2 (50U/ml) ± IL-21 (10ng/ml) with allogeneic *in vitro* generated Th1 or Th2 effectors (0.6×10^6^ cells/ml, ratio 5B:1T) for 5-6 days, as indicated. B cell media contained Iscove’s DMEM supplemented with penicillin (200μg/ml), streptomycin (200μg/ml), gentamicin (40μg/ml), 10% FBS, and insulin (5μg/ml; Santa Cruz Biotechnology).

#### B cell activation with defined stimuli

Purified B cell subsets isolated from the tonsil or blood of HD or SLE patients were cultured (1×10^6^ cells/ml) for 3 days with 5μg/ml anti-Ig (Jackson ImmunoResearch), 5μg/ml R848 (InvivoGen), 50U/ml IL-2, 10ng/ml Baff, 10ng/ml IL-21 (Peprotech) and 20ng/ml IFNγ (R&D) (Step 1). Cells were either directly analyzed or washed and recultured (2×10^5^ cells/ml) for an additional 3 days with the same stimuli (Step 2). The number of ASCs and total cells recovered in cultures on day 6 were determined and then normalized based on cell input. In some experiments, anti-Ig, R848, IL-21, IL-2, IFNγ or Baff were omitted from the cultures during Step 1, or Step 2 or both steps. In other experiments, the concentration of R848 in Step 1 and Step 2 and/or the concentration of IFNγ in Step1 was varied, as indicated in the text. In some experiments, B cell subsets isolated from blood of SLE patients and HD were stimulated for 2.5-6 days with R848 and IL-21, IL-2, Baff and IFNγ.

#### STAT3 phosphorylation assays

HD B_N_ cells were cultured with 5μg/ml anti-Ig and 5μg/ml R848 alone (Be.0) or in combination with IFNγ (Be.IFNγ), IL2 (Be.IL2), or IL2 plus IFNγ (Be.γ2) and analyzed on day 3 or were washed and recultured in the presence of IL21 and R848 for one additional day and analyzed on day 4. On day 3 or 4, cells were washed and restimulated with medium alone or with IL21 (10ng/ml) for 20 min at 37ºC. The cells were fixed and permeabilized with BD Transcription Factor Phospho Buffer Set and intracellular staining with anti phospho-STAT3 was performed.

#### *In vitro* B cell proliferation

Purified B cell subsets (1-5×10^6^ cells/ml) were stained for 10 min at 37°C with PBS diluted CellTrace™ Violet (Molecular Probes, Thermofisher). The cells were washed and either used in T effector co-culture experiments or were cultured in the presence of defined stimuli.

#### *In vitro* ASC differentiation

B_N_ cells were co-cultured with *in vitro* generated Th1 or Th2 cells plus IL-2 and IL-21. On day 6 of the co-culture B_DN_ cells from both cultures were sort-purified and then cultured in 0.22μM-filtered conditioned media (media collected from the original T/B co-cultures). ASCs were enumerated after 18 hrs by flow cytometry.

#### Cytokine measurements

Th1 and Th2 cells were restimulated with platebound anti-CD3 and anti-CD28 (both 5μg/ml). Cytokine levels in restimulated T cell cultures and SLE patient plasma samples was measured using Milliplex^®^ MAG Human Cytokine/Chemokine Immunoassays (Millipore).

#### ELISPOT

Serial diluted B cells were transferred directly to anti-IgG (Jackson ImmunoResearch) coated ELISPOT plates (Millipore) for 6 hrs. Bound Ab was detected with alkaline phosphatase-conjugated anti-human IgG (Jackson ImmunoResearch) followed by development with alkaline phosphatase substrate (Moss, Inc). ELISPOTs were visualized using a CTL ELISPOT reader. The number of spots detected per well (following correction for non-specific background) was calculated.

#### Anti-SMITH ELISAs

Anti-Smith IgG autoantibodies in plasma from SLE patients and healthy donors were detected using the enzymatic immunoassay kit (Alpha Diagnostic) according to the manufacturer protocol.

#### Flow Cytometry

Single cell suspensions were blocked with 10 μg/ml FcR blocking mAb 2.4G2 (mouse cells) or with 2% human serum or human FcR blocking reagent (Miltenyi) (human cells) and then stained with fluorochrome-conjugated Abs. 7AAD or LIVE/DEAD^®^ Fixable Dead Cell Stain Kits (Molecular Probes/ThermoFisher) were used to identify live cells. For intracellular staining, cells were stained with Abs specific for cell surface markers, fixed with formalin solution (neutral buffered, 10%; Sigma) and permeabilized with 0.1% IGEPAL (Sigma) in the presence of Abs. Alternatively, the transcription factor and phospho-transcription factor staining buffers (eBioscience) were used. Stained cells were analyzed using a FACSCanto II (BD Bioscience). Cells were sort-purified with a FACSAria (BD Biosciences) located in the UAB Comprehensive Flow Cytometry Core. Analysis was performed using FlowJo v9.9.3 and FlowJo v10.2.

#### RNA-seq library preparation and analysis

RNA samples were isolated from TRIzol (FisherThermo) treated sort-purified day 6 Be1 and Be2 IgD^neg^CD27^neg^ B cells. 300 ng of total RNA from three biological replicates per B cell subset was used as input for the KAPA stranded mRNA-seq Kit with mRNA capture beads (KAPA Biosystems). Libraries were assessed for quality on a bioanalyzer, pooled, and sequenced using 50 bp paired-end chemistry on a HiSeq2500. Sequencing reads were mapped to the hg19 version of the human genome using TopHat with the default settings and the hg19 UCSC KnownGene table as a reference transcriptome. For each gene, the overlap of reads in exons was summarized using the GenomicRanges package in R/Bioconductor. Genes that contained 2 or more reads in at least 3 samples were deemed expressed (11598 of 23056) and used as input for edgeR to identify differentially expressed genes (DEGs). *P*-values were false-discovery rate (FDR) corrected using the Benjamini-Hochberg method and genes with a FDR of <0.05 were considered significant. Expression data was normalized to reads per kilobase per million mapped reads (FPKM). Data processing and visualization scripts are available (78). All RNA-seq data is available from the GEO database under the accession GSE95282. See also Supplementary File 1.

#### ATAC-seq preparation and analysis

ATAC-seq data generated from the SLE B cell subsets was previously reported (19). ATAC-seq analysis on *in vitro* generated B cell was performed on 10,000 Be.0, Be.IFNγ, Be.IL2 or Be.γ2 cells as previously described (79, 80). Sorted cells were resuspended in 25 μl tagmentation reaction buffer (2.5 μl Tn5, 1x Tagment DNA Buffer, 0.2% Digitonin) and incubated for 1 hr at 37°C. Cells were lysed with 25 μl 2x Lysis Buffer (300 mM NaCl, 100 mM EDTA, 0.6% SDS, 1.6 μg Proteinase-K) for 30 min at 40°C, low molecular weight DNA was purified by size-selection with SPRI-beads (Agencourt), and then PCR amplified using Nextera primers with 2x HiFi Polymerase Master Mix (KAPA Biosystems). Amplified, low molecular weight DNA was isolated using a second SPRI-bead size selection. Libraries were sequenced using a 50bp paired-end run at the NYU Genome Technology Center. Raw sequencing reads were mapped to the hg19 version of the human genome using Bowtie (81) with the default settings. Duplicate reads were marked using the Picard Tools MarkDuplicates function (http://broadinstitute.github.io/picard/) and eliminated from downstream analyses. Enriched accessible peaks were identified using MACS2 (82) with the default settings. Differentially accessible regions were identified using edgeR v3.18.1 (83) and a generalized linear model. Read counts for all peaks were annotated for each sample from the bam file using the Genomic Ranges (84) R/Bioconductor package and normalized to reads per million (rpm) as previously described (80). Peaks with a greater than 2-fold change and FDR < 0.05 between comparisons were termed significant. Genomic and motif annotations were computed for ATAC-seq peaks using the HOMER (55) annotatePeaks.pl script. The findMotifsGenome.pl function of HOMER v4.8.2 (55) was used to identify motifs enriched in DAR and the ‘de novo’ output was used for downstream analysis. To generate motif footprints, the motifs occurring in peaks were annotated with the HOMER v4.8.2 annotatePeaks.pl function (55) using the options ‘-size given’. The read depth at the motif and surrounding sequence was computed using the GenomicRanges v1.22.4 (84) package and custom scripts in R/Bioconductor. All other analyses and data display were performed using R/Bioconductor with custom scripts (78). ATAC-seq data has been deposited in the NCBI GEO database under accession number GSE119726. See also Supplementary Files 3-5 for complete list of DARs and for analysis of TF motif enrichment in the ATAC-seq dataset.

#### GSEA

For gene set enrichment analysis samples were submitted to the GSEA program (http://software.broadinstitute.org/gsea/index.jsp). For the comparison of interest (i.e., B_DN_ Be1 and B_DN_ Be2 cells), all detected genes were ranked by multiplying the −log_10_ of the P-value from edgeR by the sign of the fold change and used as input for the GSEA Preranked analysis. The custom gene set defining genes upregulated in SLE T-bet^hi^ B_DN_ relative to other B cell subsets were derived from (19) and are listed in Supplementary File 2.

#### Ingenuity Pathway Analysis (IPA)

IPA upstream regulator analysis ((85), Qiagen, Redwood City CA) was performed using the log_2_ fold-change in gene expression between genes that were significantly differentially expressed (FDR < 0.05) in B_DN_ Be1 and B_DN_ Be2 cells. Upstream regulators with an activation z-score of ≥2 or ≤ −2 were considered to be activated or inhibited in B_DN_ Be1 cells. Overlap *P*-value (between the regulator’s downstream target list and the DEG list was based on Fisher’s exact test.

#### Statistical Analysis

Comparisons between two groups were performed with the Student’s *t* test for normally distributed variables and the Mann-Whitney test for non-normally distributed variables. The one-way ANOVA test was used to compare mean values of three or more groups and the Kruskal-Wallis nonparametric test was used to compare medians. Strength and direction of association between two variables measures was performed using the D’Agostino-Pearson normality test followed by Pearson’s or Spearman’s correlation test. Data were considered significant when *P* ≤ 0.05. Analysis of the data was done using the GradhPad Prism version 7.0a software (GraphPad). See Supplementary File 6 for all statistical comparisons.

#### Mice and bone marrow chimeras

All experimental animals were bred and maintained in the UAB animal facilities. All procedures involving animals were approved by the UAB Institutional Animal Care and Use Committee and were conducted in accordance with the principles outlined by the National Research Council. B6.SB-*Yaa*/J.B6;129S-*Fcgr2b*^*tm1Ttk*^/J (Yaa.*Fcgr2b*^-/-^) (75) (obtained by permission from Dr. Sylvia Bolland (NIH)) were intercrossed with B6.129S2-Ighm^tm^1^Cgn^/J (μMT) or B6.129S6-*Tbx21*^tm^1^Glm^/J (*Tbx21*^-/-^) mice (both strains obtained from Jackson Laboratory) to produce B cell deficient (Yaa.*Fcgr2b*^-/-^.μMT) or T-bet deficient (Yaa.*Fcgr2b*^-/-^.*Tbx21*^-/-^) lupus-prone mice. To generate bone marrow chimeras, μMT recipient mice were irradiated with 950 Rads from a high-energy X-ray source, delivered in a split dose 4 hrs apart. Recipients were reconstituted (10^7^ total BM cells) with 80% Yaa.*Fcgr2b*^-/-^.μMT BM + 20% Yaa.*Fcgr2b*^-/-^.*Tbx21*^-/-^ BM (B-YFT chimeras) or with 80% Yaa.*Fcgr2b*^-/-^ BM + 20% Yaa.*Fcgr2b*^-/-^.*Tbx21*^-/-^ BM (20%Control chimeras).

#### Mouse ANA detection and imaging

Antinuclear antibodies (ANA) were detected by an indirect immunofluorescence assay using HEp-2 cells. Fixed HEp-2-coated microscope slides (Kallestad®, BioRad) were blocked, incubated with serum diluted 1:100 and stained with anti-IgG-FITC (Southern Biotech) (10 μg/ml). Slides were mounted with SlowFade® Gold Antifade Mountant with DAPI (ThermoFisher) and imaged. Anti-nuclear staining was quantitated as the mean flourescence intensity (MFI) of IgG-FITC over DAPI-staining areas (nuclei) using NIS-Elements AR software (Nikon). Data are presented as log nuclear IgG MFI normalized by subtracting the MFI of negative control serum from B6 mice. ANA images were collected using a Nikon Eclipse Ti inverted microscope and recorded with a Clara interline CCD camera (Andor). The images were taken with a 20X (immunofluorescence) objective for 200-400X final magnification. Images were collected using NIS Elements software, scale bars were added and images were saved as high-resolution JPEGs. JPEG images were imported into Canvas Ver 12 software and were resized, cropped with the identical settings applied to all immunofluorescence images from the same experiment. Final images presented at 600-650 dpi (ANA).

#### Urinary Albumin to Creatinine Ratio (UACR)

Albumin concentrations in urine samples, collected from live or euthanized mice, were measured using the Mouse Albumin ELISA Quantitation Set (Bethyl Labs) according to manufacturer’s protocol using a mouse reference serum as an albumin standard. To normalize for urine concentration, urinary creatinine was measured by liquid chromatography-mass spectrometry in the UAB/UCSD O’Brien Core Center for Acute Kidney Injury Research. The UACR was calculated as μg/ml albumin divided by mg/ml creatinine and is reported as μg albumin/mg creatinine.

## Acknowledgements.

We thank Thomas Scott Simpler, Uma Mudunuru, Holly Bachus, Fen Zhou, Betty Mousseau, Enid Keyser and Dr. Ji Young Hwang for technical support; Drs. Ann Marshak-Rothstein (Univ. Massachusetts), Randall Davis (UAB) and Paul Rennert for providing mice, antibodies and cell lines and Stephanie Ledbetter, Neva Gardner, Ellen Sowell and Catrena Johnson for assistance with recruitment and consenting of healthy and vaccinated subjects. We acknowledge the Tissue Procurement Facility of the NCI-supported UAB Comprehensive Cancer Center for providing remnant tonsil tissue; the Alabama Vaccine Research Clinic, the UAB RADAR biorepository and the UAB CCTS (UL1 TR001417) for assistance in procuring human samples; the UAB Animal Resources Program Comparative Pathology Laboratory for preparation of histology slides and the UAB/UCSD O’Brien Core Center for Acute Kidney Injury Research (NIH 1P30 DK 079337) for assistance with murine urine creatinine measurements. Funding for the work was provided by the US National Institutes of Health (NIH): P01 AI078907 and R01 AI110508 (to FEL), R01 AI123733 (to JMB and CDS), U19 AI109962 (to F.E.L. and T.D.R.), P01 AI 125180 (to I.S., F.E.L., JMB and CDS) and R37AI049660-11 and U19 Autoimmunity Centers of Excellence AI110483 (to I.S.). S.L.S. was partially supported by the UAB Medical Scientist Training Program NIGMS T32GM008361; A.N. received pilot grant support from the UAB AMC21 Immunology Autoimmunity and Transplantation strategic initiative and M.I.D. received support from NIAMS K23 AR062100. The UAB CCTS informatics core (T.P.) receives support from the National Center for Advancing Translational Sciences of the National Institutes of Health under award number UL1TR001417. NIH P30 AR048311 and P30 AI027767 provided support for the UAB consolidated flow cytometry core, G20RR022807-01 provided support for the UAB Animal Resources Program X-irradiator and 5UM1CA183728 provided funding for acquisition of human tonsil tissue.

## Supplemental Figures and Files

**Figure 1-figure supplement 1.**
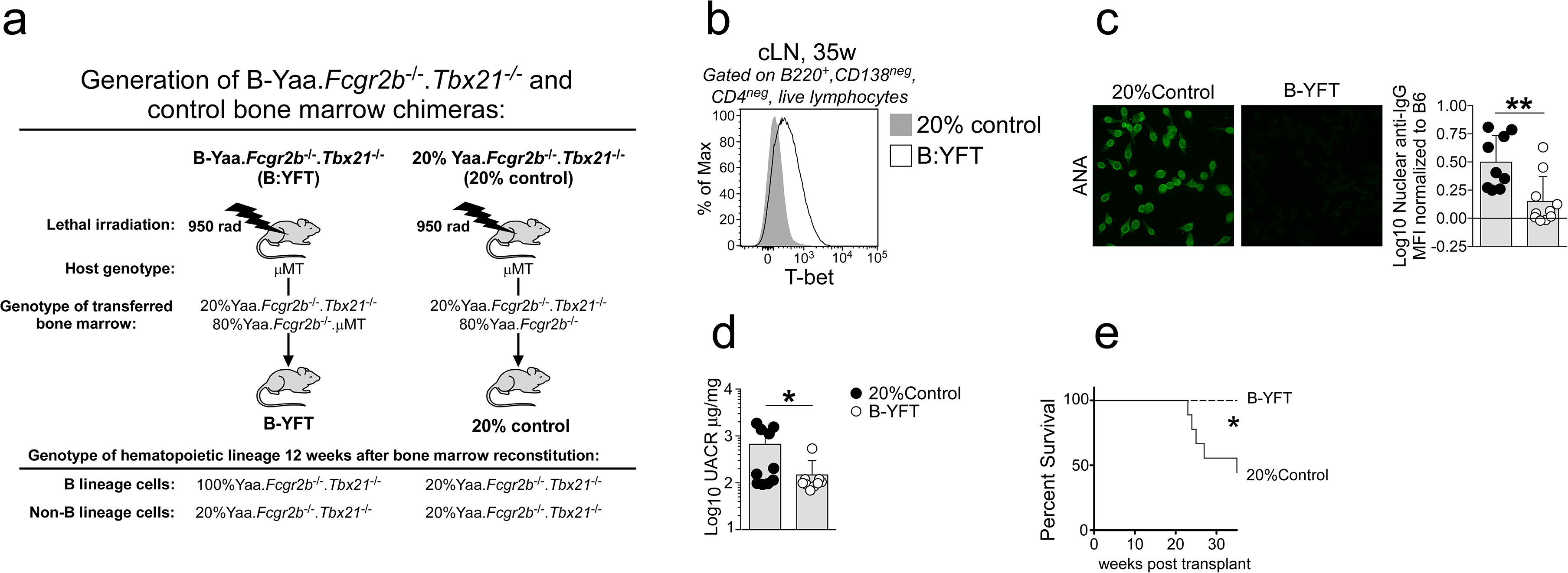
Development of SLE in TLR7-overexpressing mice requires T-bet^+^ B cells. Cartoon (**a**) depicting generation of SLE-prone bone marrow Yaa.*Fcgr2b*^-/-^ chimeras lacking T-bet in all B lineage cells or in 20% of all hematopoietic cells. To generate Yaa.*Fcgr2b*^-/-^ mice with selective deletion of T-bet in B cells (B:YFT), we reconstituted lethally irradiated B cell deficient μMT mice with a mixture of 80% B cell deficient Yaa.*Fcgr2b*^-/-^.μMT BM + 20% Yaa.*Fcgr2b*^-/-^.*Tbx21*^-/-^ BM (B-YFT chimeras). In these chimeras all hematopoietic cells, including B cells, will carry the autoimmune loci (Yaa.*Fcgr2b*^-/-^). Furthermore, <u>all B cells</u> (100%) and ∼20% of cells in all other hematopoietic cells in these animals will be T-bet deficient (*Tbx21*^-/-^). For controls (20%Control), we reconstituted irradiated μMT hosts with 80% Yaa.*Fcgr2b*^-/-^ BM + 20% Yaa.*Fcgr2b*^-/-^.*Tbx21*^-/-^ (20%Control). In these chimeras all hematopoietic cells, including B cells, will carry the autoimmune loci (Yaa.*Fcgr2b*^-/-^). In addition, 20% of all hematopoietic cells, including B cells, will be T-bet deficient. Flow cytometry analysis (**b**) showing T-bet expression by B cells isolated from the cervical lymph node (cLN) of a representative B:YFT and 20% control mouse at 35 weeks post-bone marrow reconstitution. (**c**) Representative images and quantification of anti-nuclear antibodies (ANAs) in serum from chimeras at 24 weeks post-transplant. (**d**) Kidney function reported as the urinary albumin:creatinine ratio (UACR) in individual chimeras at 24 weeks post-transplant. (**e**) Mantel-Cox survival curve of chimeras up to 35 weeks post-transplant. Representative data shown as mean ± SD from 1 of 2 independent experiments with 7-10 mice per group. Statistical analyses were performed using a Student’s t test (**c-d**) and Mantel-Cox survival test (**e**).

**Figure 3-figure supplement 1.**
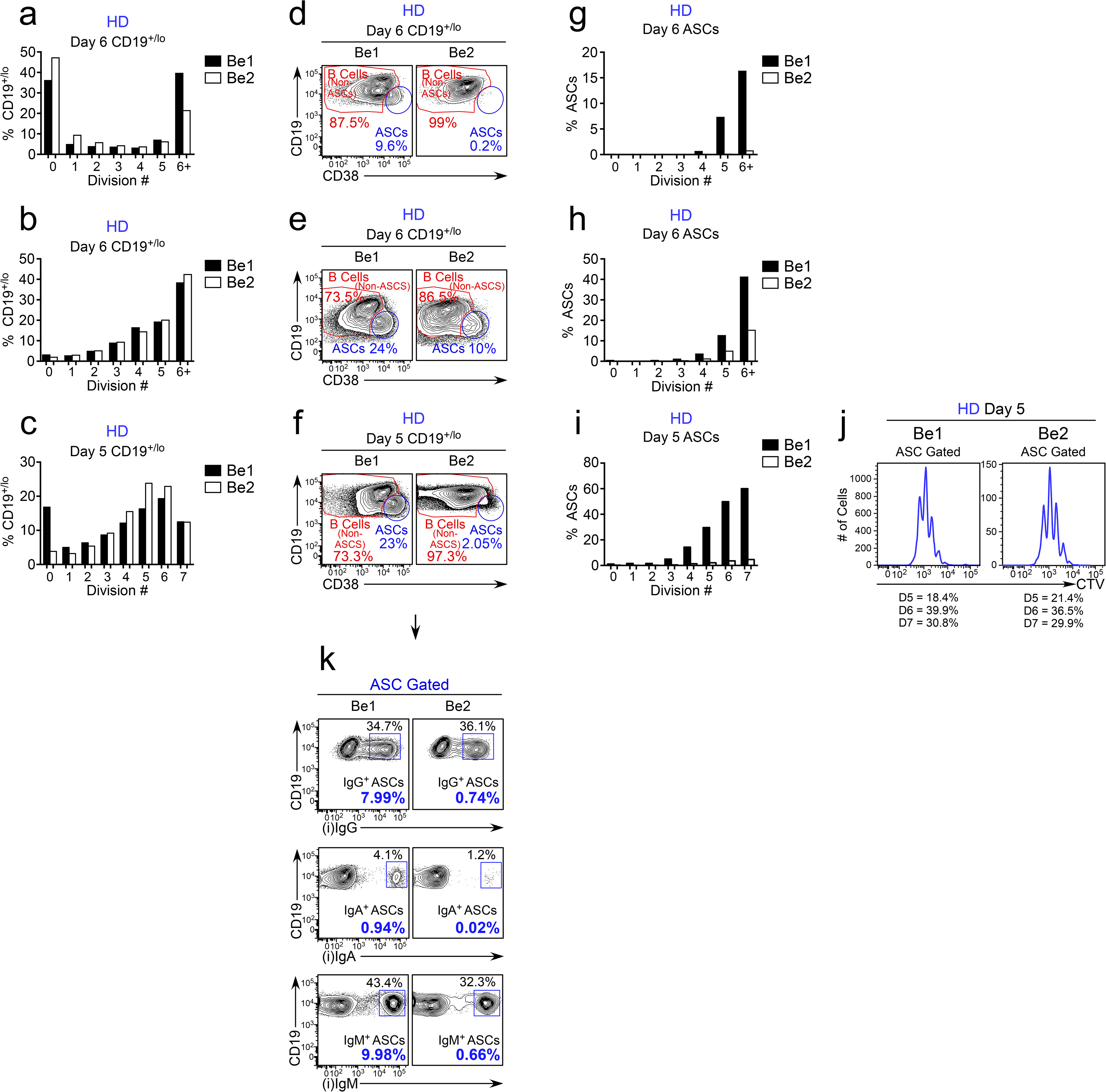
ASC formation is enhanced in co-cultures containing B_N_ and Th1 cells (Be1 co-cultures) relative to co-cultures containing B_N_ and Th2 cells (Be2 co-cultures). (**a-j**) Proliferation analysis of ASCs and B cells in 3 independent paired day 5-6 Be1 and Be2 co-cultures. Co-cultures generated with purified CTV-labeled HD B_N_ cells and allogeneic Th1 or Th2 cells + IL-21 and IL-2. B lineage cells gated as CD19^+/lo^, which includes both CD19^lo^CD38^hi^ ASCs and non-ASC B cells (see panels **d-f** for representative flow plots). Data reported as the proportion of total CD19^+/lo^ B lineage cells (**a-c**) in each cell division or the fraction of cells within each cell division that are ASCs (**g-i**). Proliferation history of ASCs in Be1 and Be2 cultures (**j**) reports the fraction of ASCs from the Be1 and Be2 co-cultures (shown in panel **c, f, i**) that are in division 5, 6 and 7. Although there is a 10-fold reduction in ASCs in Be2 co-cultures, the ASCs that are present in the Be2 co-cultures have divided the same number of times as the ASCs in the Be1 co-cultures. (**k**) Gating strategy to identify unswitched and isotype switched ASCs in Be1 and Be2 co-cultures. Representative flow plots showing intracellular IgM, IgA and IgG staining on ASC-gated cells (from panel **f**) in paired Be1 and Be2 co-cultures. Data are reported as the frequency of intracellular IgM, IgG or IgA expressing CD38^hi^CD19^lo^ ASCs within either the total CD19^+/lo^ B lineage compartment (black font) or within the total ASCs (bold purple font).

**Supplementary File 1. RNA-seq analysis of *in vitro* generated IgD^neg^CD27^neg^ B_DN_ Be1 and Be2 cells.** RNA-seq analysis of sorted IgD^neg^CD27^neg^ B_DN_ Be1 and Be2 cells isolated from Th1/B_N_ and Th2/B_N_ co-cultures. Data are shown as rpkm values from 3 independent Be1 and Be2 co-cultures that were set up with donor-matched sets of allogeneic B_N_ cells and *in vitro* polarized Th1 or Th2 cells. Log2 fold change (Be1/Be2), *P* and FDR values reported.

**Supplementary File 2. Up DEG list from T-bet expressing B_DN_ cells from SLE patients.** RNA-seq analysis was previously performed (19) on sort-purified T-bet^hi^-expressing IgD^neg^CD27^neg^IgG^+^CXCR5^neg^ B cells from HD and SLE patients (DN2 cells). The DN2 Up DEG list is defined as genes that are significantly upregulated in SLE and HD DN2 cells relative to at least one other B cell subset (B_N_, B_SW_ or CXCR5-expressing (T-bet^lo^) DN1 cells).

**Supplementary File 3. ATAC-seq data set from day 3 Be.0, Be.IFNγ, Be.IL2 and Be.γ2 B cell subsets.** HD B_N_ cells were activated for 3 days with anti-Ig and R848 alone (Be.0) or in combination with: IFNγ (Be.IFNγ), IL-2 (Be.IL2) or both IFNγ + IL-2 (Be.γ2). ATAC-seq analysis was performed on DNA isolated from each B cell subset. Table includes all identified differentially accessible regions (DAR) with fold change and FDR *q* values for each comparison. N=2 independent samples/group.

**Supplementary File 4. Transcription factor motif enrichment in ATAC-seq DAR.** To identify transcription factor binding motifs that are enriched in DARs identified in the ATAC-seq data set (See Supplementary File 3), the findMotifsGenome.pl function of HOMER v4.8.2 with the ‘de novo’ output were used for analysis. DARs analyzed are upregulated in Be.IFNγ over Be.0, Be.IL-2 over Be.0 and Be.γ2 over Be.0 (separate tabs). Table includes the list of transcription factors binding motifs, which are sorted in rank order according to *P* value.

**Supplementary File 5. P values for ATAC-seq motif enrichment comparisons.** *P* values for chromatin accessibility at transcription factor consensus DNA binding motifs (T-bet, IRF4, BLIMP1, NF-kB p65 and NF-kB REL) in ATAC-seq data. Comparisons include two-sided Student’s t-test comparisons with data from day 3 Be.0, Be.IFNγ, Be.IL2 and Be.γ2 cells.

**Supplementary File 6. Complete statistical information for all data presented in this manuscript.**

